# Burning glass effect of water droplets triggers an ER-derived calcium response in the chloroplast stroma of *Arabidopsis thaliana* leaves

**DOI:** 10.1101/2024.09.07.611817

**Authors:** Dominic Kuang, Shanna Romand, Anna S. Zvereva, Bianca Maria Orlando Marchesano, Stefano Buratti, Ke Zheng, Evelien Mylle, Cornelia Spetea, Daniël Van Damme, Bernhard Wurzinger, Markus Schwarzländer, Markus Teige, Alex Costa, Simon Stael

## Abstract

Plants require water and light for photosynthesis, but light, when focused by water droplets on leaves, can create high light intensity spots that are harmful to plants. As excessive light intensity can reduce growth or even induce cell death, it is vital for plants to detect and react to changes in light exposure and acclimate to high light stress. Ca^2+^ signaling was previously implicated in high light acclimation. However, the dynamics of free Ca^2+^ concentration in the chloroplast, the primary site of photosynthesis, or in the nucleus and in the cytoplasm, where transcription and translation for long-term acclimation occurs, remain unknown. Here we studied the dynamics and mechanism of the Ca^2+^ response to high light exposure. Focusing light through a glass bead to mimic water droplets triggered an increase of the free Ca^2+^ concentration in the chloroplast stroma of *Arabidopsis thaliana*. This finding was corroborated using established and newly developed genetically encoded calcium indicators, which revealed a biphasic increase in the stromal free Ca^2+^ concentration when exposed to varying intensities and qualities of light. Among photosynthetic by-products, reactive oxygen and lipophilic species in particular, have been implicated in high light stress acclimation. A H_2_O_2_ signature was induced, albeit with different dynamics than the Ca^2+^ response, while chemical inhibition of the photosynthetic electron transport points towards singlet oxygen as a potential culprit of the high light-induced increase in stromal free Ca^2+^ concentration. The observed dynamics differed from those of a heat-shock induced Ca^2+^ signature, although temperature had a positive effect on the Ca^2+^ response. Based on Ca^2+^ inhibitor treatments and the free Ca^2+^ concentration dynamics, we suggest that the high light-induced stromal Ca^2+^ is derived from the endoplasmic reticulum rather than from the cytoplasm. In conclusion, inspired by the burning glass effect of water droplets on leaves, we uncovered a Ca^2+^ response that implicates a novel mechanism for plants to acclimate to high light stress—a process that will become increasingly relevant in a changing climate.

## Introduction

Plants require water and light as a driving force for photosynthesis to support their development and growth. While the influence of rain and sunshine on plant growth and development has been extensively investigated in terms of water and energy availability, recent studies have begun to explore the more subtle effects of these environmental factors. For example, rain droplets can be detected through the bending of trichomes, specialized hair-like structures on the leaf surface, triggering concentric circles of calcium signals expanding outward from the base of the trichome [1]. Furthermore, water spray alone induces the production of jasmonic acid [2], a plant hormone that plays a crucial role in various physiological processes, including plant growth, development, and defense [3, 4]. These studies highlight the plant’s capacity to perceive and react to the physical disturbance that water causes to leaves.

Beyond the physical impact of rain droplets on plant physiology, the interaction of light with water on plant surfaces can create unique optical phenomena. The lensing effect, arising from the spherical shape and refractive properties of water droplets, can focus sunlight onto specific areas of the leaf. This effect is influenced by various factors, including the size, shape, and distribution of water droplets, as well as the angle of the sunlight [5]. Other potential sources of water droplets on the leaf surface can come from morning dew or guttation drops [6], a watery liquid exuded from hydathode pores at leaf edges. The surface tension of water interacting with the waxy layer of the leaf cuticle, a barrier required to control evaporation and pathogen defense [7], minimizes the surface area and determines the shape and stability of water droplets on the leaf. The combination of these factors can lead to the formation of localized regions of high light (HL) intensity on the leaf surface [5]. Understanding the mechanisms involved in the formation and effects of these localized HL zones could provide valuable insights into plant stress responses, photosynthesis, and the evolution of plant adaptations to fluctuating environmental conditions.

Exposure to excessive light intensity can be deleterious for growth. HL stress occurs when a change in light exposure exceeds the plant’s ability to manage the excess of energy that can neither be used to assimilate CO_2_ nor effectively be dissipated. This condition usually occurs in natural habitats in combination with biotic and abiotic stresses. Plants can acclimate to HL stress through short-term and long-term regulatory mechanisms [8, 9]. For example, the dissipation of excess excitation energy as heat, known as non-photochemical quenching, is induced within seconds of HL exposure [9]. The chloroplast avoidance movement, which relocates the chloroplasts along the anticlinal sides of the cells, occurs within minutes upon HL [10]. Later, light acclimation happens within hours to days as a result of complex transcriptional, translational, and post-translational gene regulation, leading to changes in the composition of the photosynthetic apparatus, anthocyanin biosynthesis and leaf morphology [8, 11, 12]. Eventually, photodamaged chloroplasts are removed by autophagy and often cell death occurs [13, 14].

HL damages mainly photosystem II and leads to the inactivation of electron transport and subsequent oxidative damage of the reaction center proteins, in particular of the D1 subunit [15]. The production of reactive oxygen species (ROS) is an integral part of the HL stress response, as overexcitation of the photosynthetic electron chain leads to production of singlet oxygen (^1^O_2_) and superoxide (O_2_ ^•−^) that is rapidly converted to hydrogen peroxide (H_2_O_2_). The highest stability of H_2_O_2_ among the various ROS makes it the most suitable second messenger of HL stress [16–18], but the highly reactive ^1^O_2_ is also implicated in HL acclimation [19, 20]. A myriad of second messengers interact and converge on retrograde signaling from the chloroplast to the nucleus for the plant to acclimate to HL stress [8].

In contrast to ROS, the potential involvement of calcium ion (Ca^2+^) signaling in the HL response is less explored. Ca^2+^ ions play essential metabolic and structural roles in chloroplasts, for example in the regulation of photosynthesis and as a component of the oxygen-evolving complex [21, 22]. Elevated Ca^2+^ concentrations in the chloroplast stroma ([Ca^2+^]_str_) were observed in response to a range of stimuli [23–26], including when plants are transferred from light to darkness [27, 28]. Several transporters and channels have been reported to mediate calcium signaling in the chloroplast: MITOCHONDRIAL CALCIUM UNIPORTER (cMCU) [26], the non-selective BIVALENT CATION TRANSPORTER 1/CA^2+^/H^+^ ANTIPORTER1/PHOTOSYNTHESIS AFFECTED MUTANT71 (BICAT1/CCHA1/PAM71) [29–31], BIVALENT CATION TRANSPORTER 2/CALCIUM/MANGANESE CATION TRANSPORTER 1 (BICAT2/CMT1)[29, 32], and the two POLLUX-like proteins PLASTID ENVELOPE ION CHANNELS 1 and 2 (PEC1/2), although it is unclear if PEC1 and PEC2 are permeable to Ca^2+^ or work in concert with yet unknown Ca^2+^-permeable channels [33]. However, none of these channels and transporters has been connected to HL acclimation. Downstream, potential targets of Ca^2+^ signaling are still unclear, although calcium binding proteins have been identified in chloroplasts [34]. Notably, CALCIUM SENSING RECEPTOR (CAS), a thylakoid-membrane-localized calcium binding protein, was found to be phosphorylated in a HL- and Ca^2+^-dependent manner and to be required for efficient acclimation to HL stress in *Arabidopsis thaliana* (hereafter Arabidopsis) [35–37]. In the single-cell green alga *Chlamydomonas reinhardtii* (hereafter Chlamydomonas), CAS directs the expression of photoprotective genes upon HL and knockdown lines are sensitive to HL [38]. A Calredoxin protein that combines four Ca^2+^-binding EF-hands with a redox-regulatory thioredoxin (TRX) was linked to antioxidant defense and HL stress in Chlamydomonas but is not conserved in plants [39, 40]. These reports implicate a potential role for stromal Ca^2+^ concentration ([Ca^2+^]_str_) dynamics in HL acclimation.

Ca^2+^ dynamics can be visualized using genetically encoded calcium indicators (GECIs). These GECIs however require light as in- and/or output (fluorescence and chemiluminescence) complicating the use of external sources of (high) light. Likely owing to these difficulties, the potential existence, dynamics, mechanism of induction, and origin of HL-induced [Ca^2+^]_str_ have remained obscure for long and were only recently observed in Chlamydomonas and the diatom *Phaeodactylum tricornutum* (hereafter Phaeodactylum) [41, 42]. Drawing inspiration from the way light is focused by water droplets creating spots of intense brightness on the leaf surface (Fig. 1A and 1B), we report the discovery of a HL-induced Ca^2+^ response in Arabidopsis. To experimentally investigate this phenomenon, we employed high-intensity lasers and white light sources to replicate the water droplet lensing effect, enabling precise control and high spatiotemporal resolution in tracking the Ca^2+^ dynamics.

**Fig. 1.**
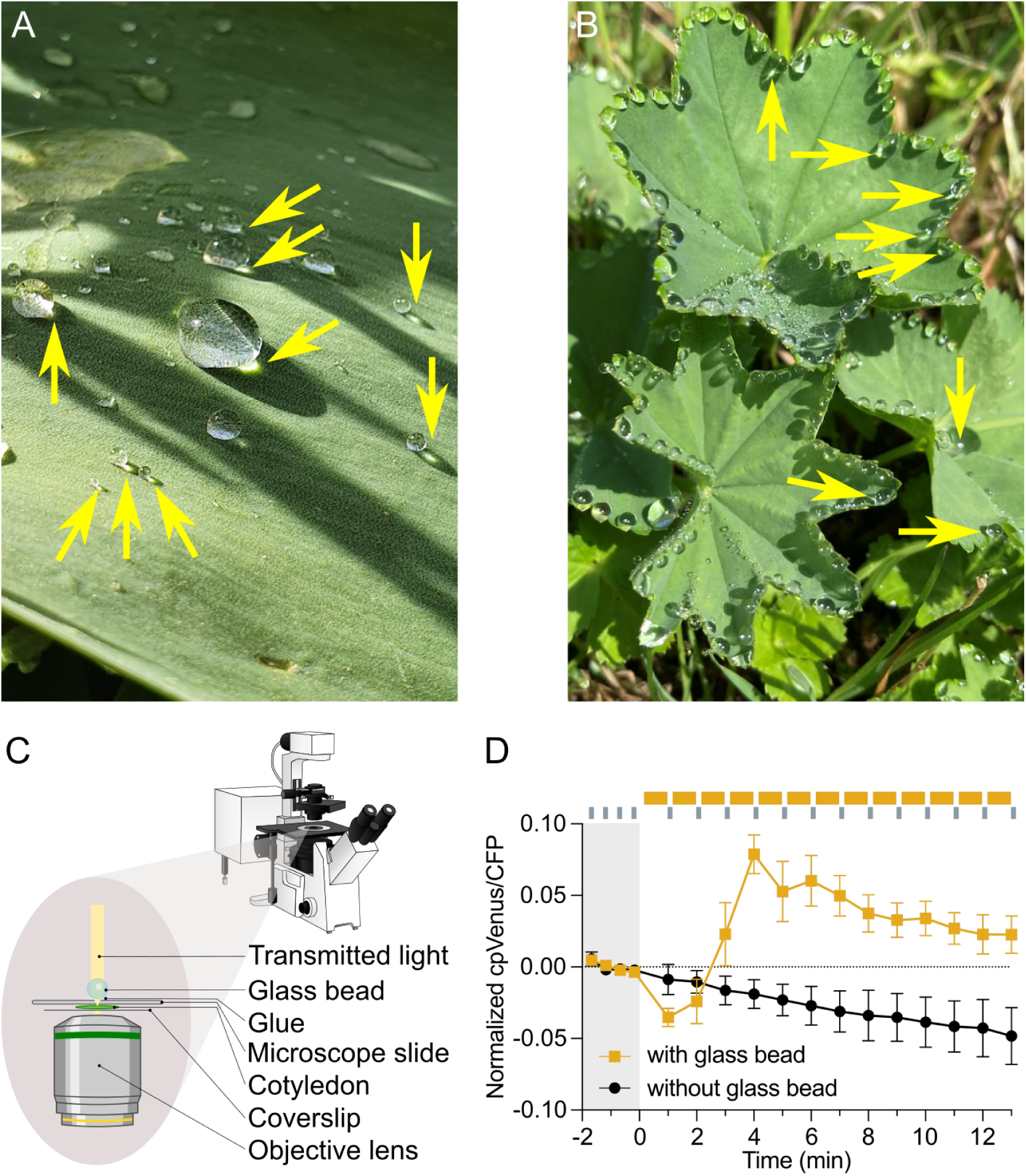
Focusing of light triggers [Ca^2+^]_str_ dynamics. **(A)** Pictures of water drops on tulip leaves (*Tulipa × gesneriana*) outside in the sun after a rain shower. **(B)** Guttation drops at the leaf edge of Ladýs Mantle (*Alchemilla sp.*). Sunlight is focused through the water drops and projected on the leaf surface (indicated with yellow arrows in (A) and (B)). **(C)** Diagram of the glass bead setup that was used to mimic the focus effect of water drops, on a confocal microscope. Transmitted light is focused through a glass bead glued to the microscope slide. **(D)** [Ca^2+^]_str_ dynamics induced by focused white halogen light (with glass bead, n = 8) and by non-focused white halogen light (without glass bead, n = 5) in cotyledons of the 2BAM4-YC3.6 expressing Arabidopsis line. Light treatment is indicated with a yellow bar. Cotyledons were intermittently imaged every 1 min between the white light stimuli. Curves represent mean ± SEM. Confocal-scanning-laser-microscope-CSLM icon by DBCLS https://togotv.dbcls.jp/en/pics.html licensed under CC-BY 4.0.

## Materials and Methods

### Arabidopsis lines and growth conditions

Arabidopsis chloroplast-localized GECI lines used in this study are NTRC-R-GECO1 (in Col-0 background) and 2BAM4-YC3.6 (in the *rdr6* background) [43]. The cytosol-localized GECI lines are R-GECO1 (in Col-0 background, a kind gift from Dr. Melanie Krebs, Heidelberg University, Germany) [44] and NES-YC3.6 (in the *rdr6* background) [43]. As a pH sensor for the chloroplast stroma we employed cp-cpYFP, and for the cytosol cyto-cpYFP (Elsässer, Feitosa-Araujo et al. 2020). As chloroplast-localized sensors for glutathione redox potential (E_GSH_) and H_2_O_2_ we selected TKTP-Grx1-roGFP2 and TKTP-roGFP2-Orp1 respectively [45], and for sensing H_2_O_2_ in the cytosol cyto-roGFP2-Orp1 [46]. The NTRC-R-GECO1 line was crossed with a cyto-nuclear localized GCaMP3 line [47] (a kind gift from Prof. Simon Gilroy, University of Wisconsin-Madison, USA, and Prof. Masatsugu Toyota, Saitama University, Japan) and stable double homozygous lines were selected.

For cotyledons/roots imaging, Arabidopsis seedlings were grown on plates containing half-strength Murashige and Skoog medium (1/2 MS) with 0.8% plant agar under long-day conditions (16h light, 110 μmol photons m^−2^ s^−1^, 22°C and 8 hours dark, 20°C) and a relative humidity of 60%. Imaging was conducted on 10- to 12-day-old seedlings.

To image soil-grown Arabidopsis plants using confocal laser scanning microscopy, NTRC-R-GECO1 seeds were sown directly in soil-filled 3.5 cm pots and kept at 4°C in the dark for 2 days to stratify. The pots were then transferred to the growth chamber under short-day conditions (12 hours light/12 hours dark, 100 µmol photons m^−2^ s^−1^) at temperatures of 18-20°C and 75% relative humidity. Imaging experiments were conducted on 7- to 8-week-old plants, when the leaves were sufficiently large to fit the customized 3D-printed chamber.

For protoplast imaging, 10- to 12-day-old seedlings were transferred from plates to soil-filled 7 cm pots and grown to maturity for 5 to 7 weeks in a chamber under long day conditions of 150 µmol photons m^−2^ s^−1^ light for 16 h, 8 h dark, 22 °C, 70% humidity.

For imaging cotyledons in the light box setup, 2BAM4-YC3.6 expressing seedlings were grown on 1/2 MS medium supplemented with 0.8% agar under long-day conditions (16 h light, 30 µmol photons m^−2^ s^−1^, 22°C and 8 hours dark, 18°C) after stratification at 4°C for two days. Plants were growing at a lower light intensity than other conditions to maximize the shift in light intensity to 600 or 1100 µmol photons m^−2^ s^−1^.

### Plasmid construction and Arabidopsis genetic transformation

NTRC-R-GECO1 and AKDE1-R-GECO1 plasmids were cloned with NEBuilder from synthetic fragments into a SpeI-ApaLI linearized vector, pGPTVII-Bar-U-RGECO1 (a kind gift from Dr. Melanie Krebs, Heidelberg University, Germany) (Suppl. Fig 2B). Plasmids were introduced into Arabidopsis (Col-0) through the floral dip method and stable single T-DNA insertion homozygous lines were selected. NTRC-R-GECO1 and AKDE1-R-GECO1 lines were further selected that have a similar expression level as the original R-GECO1 line, assessed by western blot with an anti-RFP antibody (Suppl. Fig. 2E).

### Widefield microscopy

Widefield microscopy was performed on a Nikon Eclipse Ti inverted microscope for Suppl. Fig 1A and 1B. Samples were prepared under the low light of a desktop lamp. Cotyledons of 2BAM4-YC3.6 and NES-YC3.6 lines were detached and placed in imaging solution (5 mM KCl, 10 mM MES pH 5.8, and 10 mM CaCl_2_) in a custom made imaging chamber and overlaid with a slab of phytagel to image the abaxial side. Images were acquired with a 20x 0.75 NA dry objective and excitation of 436 +/− 20 nm, and emissions around 483 +/− 20 nm (CFP) and 542 +/− 20 nm (cpVenus). Images were analyzed using the ImageJ software package [48]. Ratios of cpVenus/CFP were calculated from average intensities of full frames minus a background subtraction, normalized to the pre-stimulus intensity, and plotted over time.

### Confocal laser scanning microscopy and cell imaging of plate-grown Arabidopsis seedlings

Confocal laser scanning microscopy (CLSM) was performed on a Zeiss LSM 780 inverted confocal microscope unless stated otherwise. Images were acquired with a 20x 0.75 NA dry objective. Cotyledons and leaves from 10-12 days old Arabidopsis seedlings were detached, placed in distilled water on a microscope slide, and covered with a cover slip. For roots, the intact seedling was placed on a slide with distilled water, and covered with a cover slip with the shoot part hanging outside. To minimize later focus changes, samples were left to incubate under a desk lamp for at least 10 min before imaging. The pinhole was set at an aperture of 5 μm. The incubated samples were firstly imaged intermittently every 30 secs for 100 sec. As the exposure time for every image was 900 ms, the sample was illuminated for a relatively short period during the 30 sec interval. Hence, this period of 100 sec could be considered as a low light (not HL) treatment. Afterwards, samples were imaged every second for 600 sec. This period of 600 sec is considered as a continuous HL treatment, as the 900 ms exposure fills a relatively big part of the one second interval.

For blue and red light-induced Ca^2+^ imaging, both stroma- and cytosol-localized YC3.6 was excited with a 405 nm laser, and emission of the FRET pair CFP and cpVenus was collected at 465 – 500 nm and 525 – 560 nm, respectively. Different laser intensities of the 405 nm laser (5%, 7.5%, 10%, 20%) and of 633 nm laser (10%, 30%, 50%) were used as HL stimuli. Laser power (watt) was measured using a power meter (Thorlabs Microscope Slide Power Meter, PM400 power meter console and S170C sensor head). The corresponding light intensities (μmol photons m^−2^ s^−1^) were calculated (Suppl. table. 1). Laser power was measured periodically to ensure similar light intensity, as laser strength decays over time. Both stroma/cytosol-localized R-GECO1 was excited with a 1% 561 nm laser, emission was collected at 580 – 630 nm. Along with the excitation light, a 10% 405 nm or a 30% 633 nm laser was used as HL stimulus. The imaging protocols of R-GECO1 and YC3.6 were also used on Col-0 plants, to exclude the potential influence of chlorophyll autofluorescence. For blue light-induced ER Ca^2+^ imaging, ER-localized ER-GCaMP6-210, was sequentially excited by a 10% 405 nm laser and a 20% 488 nm laser, emission was collected at 505 – 560 nm [49, 50]; for red light-induced ER Ca^2+^ imaging, ER-GCaMP6-210 was excited by a 2% 488 nm laser, with an additional constant 30% 633 nm laser as the HL stimulus. Cytosol-localized GCaMP3 of the NTRC-R-GECO1 × cyto-GCaMP3 line was excited with a 1% 488 nm laser, emission was collected at 500 – 550 nm. R-GECO1 was imaged as mentioned above with an additional 10% 405 nm laser as the HL stimulus. Chlorophyll autofluorescence was collected at an emission of 650 – 720 nm.

For blue light-induced ROS imaging, cotyledons of TKTP-roGFP2-Orp1 and cyt-roGFP2-Orp1 expressing plants were imaged with the same light regime as 2BAM4-YC3.6 for a correct comparison in Fig. 4. Cotyledons were excited sequentially with a 405 nm diode laser and a 488 nm white light laser on a Leica SP8 CSLM, with a 40x water immersion objective. For roGFP2-Orp1, emission was collected at 500-527 nm in both sequences, which is required for ratiometric imaging of roGFP2 oxidation/reduction. For YC3.6, 455-483 nm (CFP) and 514-542 nm (cpVenus) emission was collected in the 405 nm excitation sequence, required for ratiometric Ca^2+^ imaging, and emission was set out of range for the 488 nm excitation, because this excitation was needed to have an equal input of laser strength as roGFP2-Orp1 and not for imaging. Two rounds of experiments were carried out, first with 9,94% of 405 nm and 11% of 488 nm excitation, and secondly with 20% 405 nm and 22% 488 nm excitation. Cotyledons of stroma-localized glutathione redox potential (E_GSH_) sensor, TKTP-Grx1-roGFP2, were sequentially excited with a 405 nm and a 488 nm laser (energy equals to 10% 405 nm laser), emission was collected at 510-535 nm on the Zeiss LSM 780 (Suppl. Fig. 7).

Stromal pH was measured as a control for the stroma-localized R-GECO1 reporter with a cp-cpYFP reporter on the Zeiss LSM 780 (Suppl. Fig. 3B). cpYFP was sequentially excited with a 405 nm and a 488 nm laser (energy equals to 10% 405 nm laser), emission was collected at 510-535 nm.

For quantification, average intensities of full frames were calculated with ImageJ, data were normalized to the average baseline intensity, and plotted over time.

### Quantification of tissue-, chloroplast, or nuclei-specific R-GECO1 and YC3.6 signals

Regions of interest (ROIs) were drawn around groups of chloroplasts corresponding to a specific tissue (mesophyll, epidermis, and stomata) based on their morphological differences, or around single chloroplasts, using ImageJ. ROIs were drawn around single nuclei for cyt-R-GECO1 based on their typical shape (see Suppl. Fig. 9A) and behavior (lack of movement or cytoplasmic streaming). Average signal intensity per ROI was measured over time for R-GECO1 and YC3.6 and was normalized by dividing all values per time series by the average of values of the baseline. To quantify the effect of different laser strengths, the parameters ‘area under the curv’ (AUC) and ‘time to reach max’ (TRM) were calculated.

### 3D design and 3D printing

To image soil-grown Arabidopsis plants with an upright CLSM, we designed a dedicated 3D-printed chamber. The chamber was designed using the Fusion 360 3D CAD software (Autodesk, https://www.autodesk.com/). The 3D models were exported in standard tessellation language (STL) format and post-processed with the open-source slicing engine Cura (Ultimaker, https://ultimaker.com/software/ultimaker-cura). Chambers were printed with a Kobra 3D printer (Anycubic, https://store.anycubic.com/products/kobra) at 0.2 mm resolution and 100% infill in PETG standard plastic filament.

The chamber was designed for single-cell imaging of soil-grown Arabidopsis leaves while keeping them attached to the plants in their pots. To achieve this, the chamber was designed to keep the imaged leaf as flat as possible, a crucial requirement for using the Water Dipping Objective (Nikon CFI75 Apo 25XC W 1300) to focus on single cells. This was accomplished by designing a pot casing with a support similar to a “trampoline” to hold the leaf. To prevent damage to the leaf lamina, the surfaces of the “trampoline” that contact the leaf were covered with cotton strips. When short-day-grown Arabidopsis plants in their pots were transferred to the imaging chamber, a large rosette leaf was secured to the holder with two braces and placed on the microscopy stage under the objective (Suppl. Fig. 4A).

### Confocal laser scanning microscopy and chloroplasts imaging of soil-grown Arabidopsis plants

CLSM analyses were performed using a Nikon A1R laser scanning confocal mounted on a Nikon ECLIPSE FN1 upright microscope with a Water Dipping Objective (Nikon CFI75 Apo 25XC W 1300). Images were acquired at 512 x 512 pixels with 5X digital zoom. NTRC-R-GECO1 was excited by a 561 nm single-mode optical fiber laser, and the emission was collected at 580-630 nm. The 561 nm laser line was set to 1%, corresponding to an incident power of 26 μW measured on the sample with the 25X objective (Supporting Information Table S1).

Different experiments were conducted. As a control, plants expressing NTRC-R-GECO1 were illuminated with the 561 nm laser set to 1% every 5 secs for 16 min. For HL stress experiments, two conditions were tested: i) plants expressing NTRC-R-GECO1 were illuminated with the 561 nm laser set to 1% every 5 secs for 3 min, followed by 3 min of continuous illumination, and then 10 min of illumination and acquisition every 5 secs; ii) plants expressing NTRC-R-GECO1 were illuminated with the 561 nm laser set to 1% every 5 secs for 3 min, followed by 3 min of continuous illumination with the 561 nm set to 1% laser coupled with 405 nm illumination (set to 20%, corresponding to 115-120 μW), and then 10 min of illumination with the sole 561 nm laser set to 1% and acquisition every 5 secs.

Images were analyzed using IMAGEJ software, and pixel intensities were measured over ROIs drawn on single chloroplasts. NTRC-RGECO1 emissions were used for calculations, normalized to the pre-stimulus intensity, and plotted against time. Images in Suppl. Fig. 4 and Suppl. Movie 3 were denoised using the NIS-Element Denoise.ai plugin (https://www.microscope.healthcare.nikon.com/en_EU/products/confocal-microscopes/a1hd25-a1rhd25/nis-elements-ai). Only non-denoised images were analyzed for fluorescence quantification using Fiji [48].

### Manual white light illumination and confocal laser scanning microscopy

For white light-induced Ca^2+^ imaging, the epifluorescence light source of the Zeiss LSM780 CLSM, a mercury short-arc lamp (HXP 120 V), was used to expose the cotyledons of 2BAM4-YC3.6 expressing plants to white light (Fig. 3I). For Fig. 1, the transmitted light, a Zeiss HAL 100 Halogen lamp, was used as a source of white light on the Zeiss LSM780 CLSM. To lens the transmitted light, a glass bead of 5 mm diameter was glued to the microscope slide on top of the position of the mounted cotyledon. Images were taken each minute by switching manually between white light (around 50 sec) and acquisition mode (around 10 sec). Images were taken and data processed as explained previously for plate-grown Arabidopsis seedlings.

For Suppl. Fig. 5, the transmitted light, a Zeiss HAL 100 Halogen lamp, was used as a source of white light on a Zeiss LSM710 inverted confocal microscope with a C-Apochromat 40x/1.20 W Korr M27 objective. Cotyledons of 2BAM4-YC3.6 and NES-YC3.6 plants were switched from low light (10% transmitted light) to HL (100% transmitted light). Images were taken each minute by switching manually between white light (around 50 sec) and acquisition mode (around 10 sec) for 2BAM4-YC3.6 and each half a minute for NES-YC3.6. YC3.6 was excited with a 5% 405 nm laser, and emission of FRET pair proteins CFP and cpVenus was collected at 465-500 nm and 525-560 nm, respectively. For data in Suppl. Fig 5 we performed plastid segmentation and tracking with Fiji and ilastik. Raw image files were opened in Fiji [48] and the chlorophyll autofluorescence channel only was selected for each time series using the slice keeper tool. The chlorophyll autofluorescence image series were saved in hdf5 format, without compression and in xyt dimensions, using the hdf5 image plugin from Fiji. The chlorophyll autofluorescence image series was then loaded into ilastik v1.4.0 [51] as input data and pixel classification was started at the first image of each time series. For training, two labels were created: one for the background, one for chloroplasts. Thresholding was chosen that only non-overlapping plastids remained in the selection. This eliminated about half of the mesophyll plastid signals since chloroplast crowding therein leads to merged objects but at the same time allows for more accurate tracking of the remaining single plastids. The tracking workflow was started by importing the series of images and the results of the pixel classification workflow. Following the “animal tracking workflow”, chloroplasts were semi automatically tracked through the image series. First ∼ 30 different chloroplasts were semi automatically tracked throughout a data set and then automatic tracking was used to follow the rest of the plastids throughout the datasets. The resulting object circumferences were used as masks to create ROIs on the CFP and cpVenus fluorescence channels in Fiji. Average fluorescence ratios for multiple single chloroplasts were plotted over time.

### On-stage white light illumination system and confocal laser scanning microscopy

The 2BAM4-YC3.6 line was imaged during low light-to-high light transitions using a Zeiss LSM 980 microscope equipped with a 25X lens (Plan-Apochromat) (Carl Zeiss Microscopy GmbH, Jena, Germany). Cotyledons of 15-day old seedlings were cut and mounted between two coverslips (18 x 18 mm and 22 x 40 mm, VWR International GmbH, Darmstadt, Germany). The light box setup was previously described in detail in Elsässer et al. [52]. Shortly, a customized on-stage illumination system was connected to the LSM 980 via the Zeiss trigger interface, enabling control of the microscope and the illumination system in a coordinated manner using the Experiment Designer in ZEN blue 3.5. A custom-built device (workshop, Institute for Geoinformatics, University of Münster, Germany) uses a 5 V trigger signal to switch a cold-white LED (Optonica GmbH, Salzburg, Germany) implemented in the on-stage illumination system. Images (1.26 sec per scan) were taken automatically with a 30 sec interval during which the sample is exposed to LED light. YC3.6 fluorescence was excited at 445 nm, while emission was collected at 473-490 nm (CFP) and 535-552 nm (cpVenus). The pinhole was set to 600 µm. The CLSM time series datasets were processed with a custom MATLAB-based software110 using x,y noise filtering {Fricker, M. D., 2016}. Ratios were then log10-transformed before statistical analysis was performed using GraphPad Prism (version 8.0.1, GraphPad Software, San Diego, CA, USA).

### Confocal laser scanning microscopy and chloroplast imaging of protoplasts with chemical treatments

Protoplasts were isolated from 5- to 6-week-old plants using the “Tape-Arabidopsis Sandwich” method [53]. Protoplasts isolated from the NTRC-R-GECO1 line were imaged on a Zeiss LSM 780 microscope with a 20X 0.75 NA dry objective. R-GECO1 was excited with a 1% 561 nm laser, emission was collected at 580-630 nm. A mercury short-arc lamp (HXP 120 V) was used for white high light stimuli. Protoplasts were firstly imaged every 30 sec without white light (baseline), afterwards, images were taken each minute by switching manually between white light (50 sec) and acquisition mode (10 sec). Chemical treatments on protoplast system: 10 µM 3-(3,4-dichlorophenyl)-1,1-dimethylurea (DCMU), 10 µM 2,5-Dibromo-6-isopropyl-3-methyl-1,4-benzoquinone (DBMIB), 50 μM methyl viologen (MV), 25 μM Cyclopiazonic acid (CPA), 10 mM ethylene glycol tetraacetic acid (EGTA). Protoplasts were kept in W5 solution (154 mM NaCl, 125 mM CaCl_2_, 5 mM KCl, 5 mM glucose, and 2 mM MES, pH 5.7) for all measurements, except in combination with EGTA, where we used MMg solution (0.4 M mannitol, 15 mM MgCl2, and 4 mM MES, pH 5.7). For quantification, average intensities of full frames were calculated with Fiji, data were normalized to the average baseline intensity, and plotted over time.

Protoplasts isolated from Col-0 plants were incubated with 100 µM Singlet Oxygen Sensor Green (SOSG) for 90 min. SOSG was excited with a 1% 488 nm laser, emission was collected at 500-600 nm on the Zeiss LSM 780 with a 20X 0.75 NA dry objective. The white light treatment was applied as mentioned above for NTRC-R-GECO1 protoplasts. Protoplasts without SOSG incubation were imaged using the same protocol to exclude the effect of chlorophyll autofluorescence. For quantification, average intensities of selected protoplasts were calculated with Fiji. Data were divided by background intensities to exclude the effect of SOSG photosensitization, normalized to the average baseline values, and plotted over time.

Mitochondria of protoplasts were stained with the MitoView™ 405 dye (Biotium, ref. #70070) at a concentration of 200 nM. The protoplasts were incubating for 15 min before imaging.

### Confocal spinning disk microscopy and temperature controlled imaging

Confocal spinning disk microscopy (CSDM) was performed on a Nikon Ti microscope with the Perfect Focus System (PFSIII) for Z-drift compensation, equipped with an Ultraview spinning-disk system (PerkinElmer) and two 512×512 Hamamatsu ImagEM C9100-13 EMccd cameras. Images were acquired using a 20X dry objective. Stromal Ca^2+^ imaging for 2BAM4-YC3.6 was carried out similar to the protocol mentioned above and the imaging setup is shown in Suppl. Fig. 8A. YC3.6 was excited at 10% 405 nm (149 µW), emission was passed through a 509 nm beam splitter and collected for CFP channel at 455-509 nm and cpVenus channel at 525-575 nm. Different temperature incubations (constant 10, 20, 30 or 40℃), temperature shifts (2-minute 20℃ to 13-minute 40℃ to 5-minute 20℃), and a constant 20℃ during light shifts were performed with a CherryTemp heating/cooling system (Cherry Biotech, Montreuil, France). For the temperature incubation (Fig. 5A), the light regime was switched from 2-minute intermittent imaging (once every 30 sec) to 13-minute continuous imaging; for temperature shifts / controls (Fig. 5B; Suppl. Fig. 8B, C), the light regime was 20-minute intermittent imaging (once every 10 sec); for light shifts (Fig. 5C), the light regime was switched from 2-minute intermittent imaging (once every 30 sec) to 13-minute continuous imaging to 5-minute intermittent imaging (once every 30 sec). The samples were incubated for 5 min on the CherryTemp system prior to imaging to stabilize the initial temperature.

### Protein extraction and immunoblotting

Protein extraction and immunoblotting were performed on 25 bulked seedlings. Total protein extraction was performed with Laemmli extraction buffer (0.0625 M Tris base; 0.07 M sodium dodecyl sulfate; 10% glycerol; pH 6.8) and samples were heated for 5 min at 85°C. Proteins were then reduced with 5% β-mercapto-ethanol, equal quantities separated by SDS-PAGE with precasted Mini-PROTEAN TGX Stain-Free Gels, and transferred onto PVDF membranes with the Trans-Blot Turbo Transfer System, according to the manufacturer’s instructions (Bio-Rad). Total protein normalization was confirmed by Ponceau Red staining, based on the level of the large subunit of Rubisco. After incubation with 5% (w/v) nonfat milk in PBST solution (0.14 M NaCl; 0.0027 M KCl; 0.01 M PO_4_^3-^; 0.05% Tween; pH 7.4) for 60 min, the membrane was incubated in the same solution with RFP-tag primary antibody (GenScript, ref. A00682) at a 1/200 dilution for 1h at room temperature. The membrane was then washed three times for 5 min in PBST and incubated in 5% nonfat milk in PBST with horseradish peroxidase-conjugated anti-Rabbit (Agrisera, ref. AS09 602) antibody at a 1/10 000 dilution for 1h at room temperature. The membrane was then washed a further three times in PBST, developed using Clarity Max Western ECL Substrate, and imaged with a Chemidoc XRS+ System (Bio-Rad).

### Statistical analyses

One-way ANOVA was applied at significant level α = 0.05. When there were statistically significant differences, Fisher test was applied for mean comparisons. Error bars indicate SEM (standard error of means). The numbers of independent biological replicates used to calculate SEM are indicated in the figure legends.

## Results

### Blue light elicits a biphasic Ca^2+^ response in chloroplasts

We attempted to elicit a [Ca^2+^]_str_ response by exposing cotyledons to 100 % of the transmitted light of a microscope and intermittently taking images of the fluorescent chloroplast-targeted Ca^2+^ reporter, 2BAM4-YC3.6 [43]. When this proved unsuccessful, we took inspiration from the way water drops can focus light (Fig. 1A and 1B). Indeed, we found that increasing the light intensity by focusing through a glass bead glued to the microscopy slide could trigger a [Ca^2+^]_str_ response, whereas the response was absent without the glass bead (Fig. 1C and 1D).

Historically, light-to-dark-induced [Ca^2+^]_str_ was measured with luminescent Ca^2+^ sensors such as aequorin [27], likely because of availability of these sensors at the time and the practical reason that they require darkness for their readout. We had attempted to confirm this dark-induced [Ca^2+^]_str_ increase with plants expressing 2BAM4-YC3.6. However, we realized that the blue excitation light (436 +/− 20 nm wavelength) required for imaging the 2BAM4-YC3.6 reporter [43] can trigger a [Ca^2+^]_str_ increase depending on the sampling time. When images with a duration of 300 ms were made every 60 sec, no perturbation was recorded. However, imaging with a 5 sec interval led to a [Ca^2+^]_str_ increase after a short initial lag phase (Suppl. Fig. 1A). A 90 sec pulse of blue light was sufficient to trigger the [Ca^2+^]_str_ response (Suppl. Fig 1B). In theory, altering the number of images taken during the time course of a dark-induced [Ca^2+^]_str_ increase should not influence the Ca^2+^ signal itself. Intrigued by this and coupled to the observation that light focused through a glass bead triggered a [Ca^2+^]_str_ increase (Fig. 1C and 1D), we set out to analyze this response in more detail.

To confirm that exposure to blue light triggers the elevation of [Ca^2+^]_str_, 2BAM4-YC3.6 signals were measured in the plastids of cotyledons, leaves and roots with a blue laser (405 nm). After a switch from quasi-darkness (blue light exposure for 900 ms every 30 s) to continuous illumination, a [Ca^2+^]_str_ spike occurred in cotyledons within seconds and lasted for about 30 sec (Fig. 2A and Suppl. Fig 1F). Afterwards, [Ca^2+^]_str_ returned to a basal level for 1-2 min before rising again more slowly and persistently (peak on average at 11.3 min ± 0.3 standard error of means (SEM)) (Fig. 2A). The initial [Ca^2+^]_str_ spike arose from a handful of random chloroplasts in the images (shown in Fig. 2C at 8, 16 and 40 sec). In contrast, the later persistent [Ca^2+^]_str_ increase occurred in almost all chloroplasts (Fig. 2C at 650 sec).

**Fig. 2.**
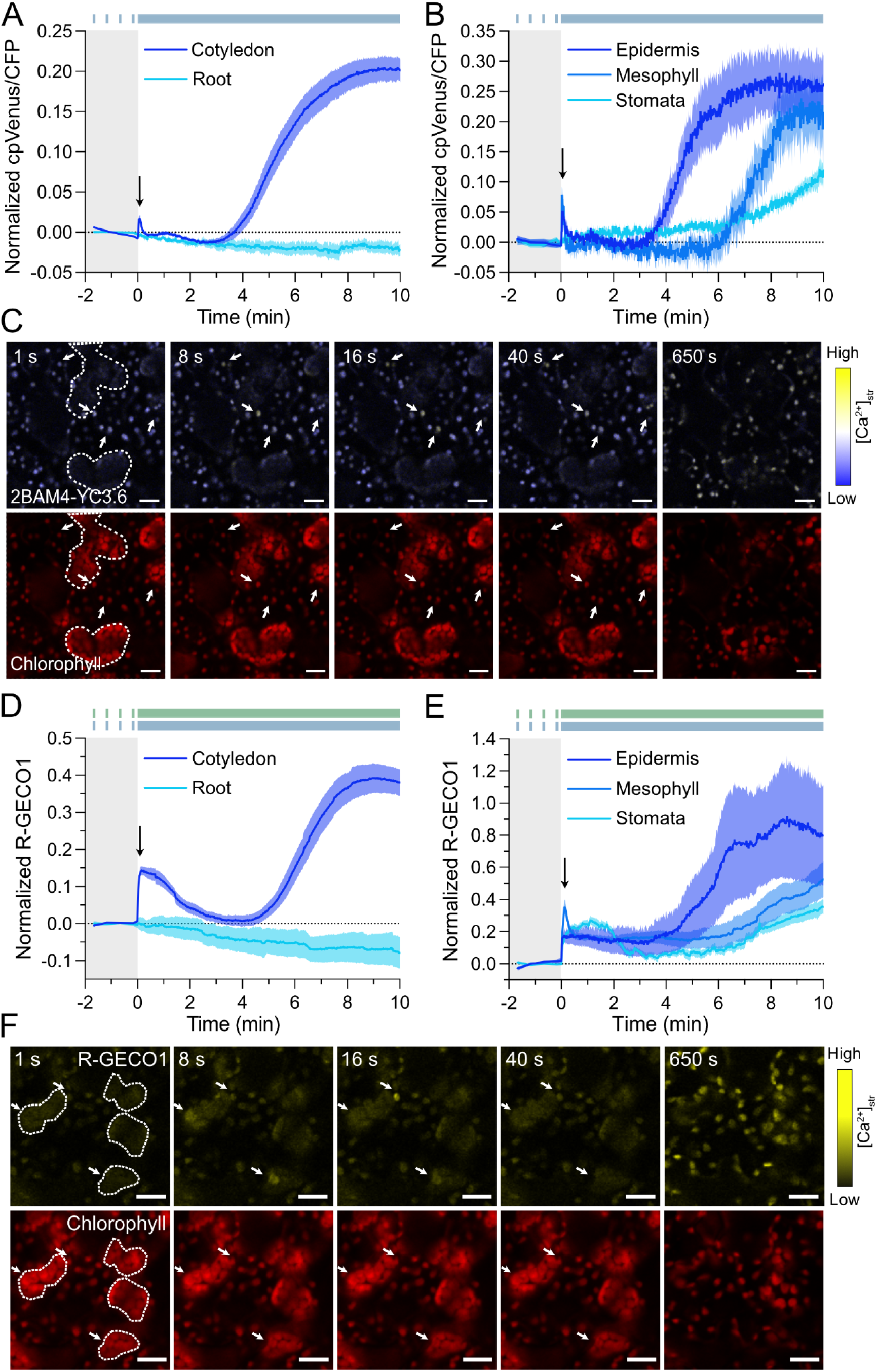
Blue light induces biphasic [Ca^2+^]_str_ dynamics in chloroplasts. **(A)** [Ca^2+^]_str_ dynamics induced by 10% 405 nm laser in cotyledons and roots of 2BAM4-YC3.6 line. n = 6. The black arrow indicates the initial fast spike of [Ca^2+^]_str_. The blue bar indicates the light regime switching from intermittent imaging (baseline) to continuous imaging (blue light stimulus). (**B**) Epidermis (n = 5), mesophyll (n = 6) and stomata (n = 12) cells of 2BAM4-YC3.6 expressing cotyledons show a heterogeneous response in [Ca^2+^]_str_ dynamics induced by 10% 405 nm laser. (**C**) Merged cpVenus and CFP signals (top) and chlorophyll (bottom) of the 2BAM4-YC3.6 expressing line. Scale bar, 20 μm. (**D**) [Ca^2+^]_str_ dynamics induced by 10% 405 nm laser in cotyledons and roots of NTRC-R-GECO1 line. NTRC-R-GECO1 was additionally excited by a 1% 561 nm laser (green bar). n = 5. (**E**) Epidermis (n = 6), mesophyll (n = 15) and stomata (n = 15) cells of cotyledons expressing NTRC-R-GECO1 show heterogeneous [Ca^2+^]_str_ kinetics induced by 10% 405 nm laser. (**F**) R-GECO1 signal (top) and chlorophyll (bottom) of NTRC-R-GECO1 line. Lines represent mean ± SEM. Scale bar, 50 μm. White arrows in 1s, 8s, 16s, 40s indicate chloroplasts with increased ratio, and white dashed lines indicate chloroplasts of mesophyll cells in (C) and (F).

Upon closer inspection, chloroplasts were found to react differently to the light stimulus depending on the tissue in which they were present (Fig. 2B). Mesophyll chloroplasts, with their characteristic large and clustered appearance, displayed a markedly delayed persistent [Ca^2+^]_str_ increase compared to the smaller and scattered epidermal chloroplasts [54] in the same field of view (Fig. 2B, C). Guard cell chloroplasts were even more delayed in their response (Fig. 2B).

First true leaves behaved similar to cotyledons. An initial spike and later increase of [Ca^2+^]_str_ were also observed upon continuous laser scanning (Suppl Fig. 1C), although the persistent [Ca^2+^]_str_ increase was delayed compared to that in cotyledons (Fig. 2A). Neither the initial spike nor the later persistent [Ca^2+^]_str_ increase could be observed in root plastids (Fig. 2A), indicating that only photosynthetically active plastids (chloroplasts) respond to continuous blue light exposure.

### A newly-established, red-shifted, and chloroplast-localized GECI reporter, NTRC-R-GECO1, confirms the blue light-induced stromal Ca^2+^ response

To gain flexibility for imaging multiple fluorescent sensors in parallel and to make our analyses more robust, we targeted the red-shifted fluorescent Ca^2+^ reporter R-GECO1 [55] to the chloroplast using the chloroplast targeting peptide of NADPH thioredoxin reductase C (NTRC-R-GECO1; Suppl. Fig. 2A and 2B). The selected lines did not show obvious differential growth phenotypes as compared to wild type (Suppl. Fig. 2C) and the NTRC-R-GECO1 signal overlapped with chlorophyll autofluorescence (Suppl Fig. 2D). NTRC-R-GECO1 allowed us to image the [Ca^2+^]_str_ response at a different wavelength (561 nm) than the blue light stimulus (405 nm). Upon continuous blue light exposure, the first [Ca^2+^]_str_ spike increased similarly fast as with 2BAM4-YC3.6, whereas it decreased more slowly (Fig. 2D). Afterwards, the persistent [Ca^2+^]_str_ transient was observed in accordance with the 2BAM4-YC3.6 experiment.

In parallel with the NTRC-R-GECO1 line, we produced a mitochondria-targeted R-GECO1 making use of the mitochondria-localized 2-oxoglutarate dehydrogenase E1 targeting peptide (AKDE1-R-GECO1; Suppl. Fig. 2). The selected lines had a similar level of R-GECO1 protein accumulation on western blot compared to NTRC-R-GECO1 and a previously established R-GECO1 line (Suppl. Fig. 2E), did not show obvious growth phenotypes compared to wild type (Suppl. Fig. 2C), and AKDE1-R-GECO1 signal overlapped with a mitochondrial dye in protoplasts (Suppl Fig. 2D). The AKDE1-R-GECO1 signal decreased over time in response to a blue light stimulus, possibly due to bleaching of the fluorophore. This indicates that bleaching likely takes place when imaging NTRC-R-GECO1, but the Ca^2+^ response is markedly different between chloroplasts and mitochondria (Suppl. Fig. 2F).

The initial [Ca^2+^]_str_ spike could also be observed when imaged only with a 561 nm laser (the higher value compared with those in Fig. 2D is due to normalization), but the later persistent [Ca^2+^]_str_ transient was missing (Suppl. Fig. 3A). R-GECO1 is known to be pH-sensitive [44]. Therefore, we estimated pH changes in the stroma with the pH sensitive reporter cpYFP [56]. Due to limitations of the imaging settings, 3.5% 561 nm laser was used as a replacement of 10% 405, as they were measured to have similar light intensities (Suppl. Table 1). Upon continuous blue light exposure, the stromal pH first increased for two min and then decreased (Suppl. Fig. 3B), similar to the NTRC-R-GECO1 signal (Fig. 2D). Consequently, the slow decrease after the initial [Ca^2+^]_str_ spike is likely explained by a pH effect on the NTRC-R-GECO1 sensor. Nevertheless, during the initial [Ca^2+^]_str_ spike, some chloroplasts in the NTRC-R-GECO1 line exhibited a stronger response than the average increase in signal (Suppl. Fig. 3C and 3D). Hence, most chloroplasts undergo stromal alkalinization as a result of the light stimulus [52] corresponding to an overall NTRC-R-GECO1 signal increase, but only a few chloroplasts experience a [Ca^2+^]_str_ spike accompanied by a larger increase of the NTRC-R-GECO1 signal. The later persistent increase of NTRC-R-GECO1 signal was exclusively due to a [Ca^2+^]_str_ increase. Root plastids of NTRC-R-GECO1 did not respond to blue light exposure (Fig. 2A, D). Epidermal and mesophyll chloroplasts also displayed different kinetics of persistent [Ca^2+^]_str_ elevations in the NTRC-R-GECO1 line, while the signatures of mesophyll and stomata chloroplasts were quite similar (Fig. 2E). This was more evident when compared with the response in the 2BAM4-YC3.6 line, likely because of a higher and more uniform expression of the NTRC-R-GECO1 reporter and its larger dynamic range (Fig. 2D, E). The initial [Ca^2+^]_str_ spike occurred in only a few seemingly random chloroplasts irrespective of tissue localization (Fig. 2F). In conclusion, the NTRC-R-GECO1 experiments faithfully recapitulated the 2BAM4-YC3.6 data.

Finally, taking advantage of a newly established setup that allows imaging soil-grown plants with confocal microscopy, we proceeded with the characterization of the blue light-induced [Ca^2+^]_str_ response in non-detached mature leaves of Arabidopsis plants expressing NTRC-R-GECO1 (Suppl. Fig. 4A and 4B). As controls, intermittent imaging with a 561 nm laser did not alter the [Ca^2+^]_str_ response (Suppl. Fig. 4C), whereas continuous imaging with the 561 nm increased the RGECO1 signal (Suppl. Fig. 4D), likely because of the effect of stromal alkalinization as was noticed in experiments with detached cotyledons (Suppl. Fig. 3A and 3B). Concomitant exposure of the leaf to a 405 nm laser produced a similar strong persistent [Ca^2+^]_str_ increase as was observed in detached cotyledon (Suppl. Fig. 3E and 3F), albeit with a faster dynamic. This could be due to the slightly stronger light stimulus (Suppl. Table 1) or an inherent faster response of non-detached mature leaves. Cotyledons and first true leaves were imaged from the abaxial side. Mature leaves, however, were imaged from the adaxial side. Hence, epidermal and mesophyll chloroplasts from both sides of the leaf reacted to the blue light stimulus.

**Fig. 3.**
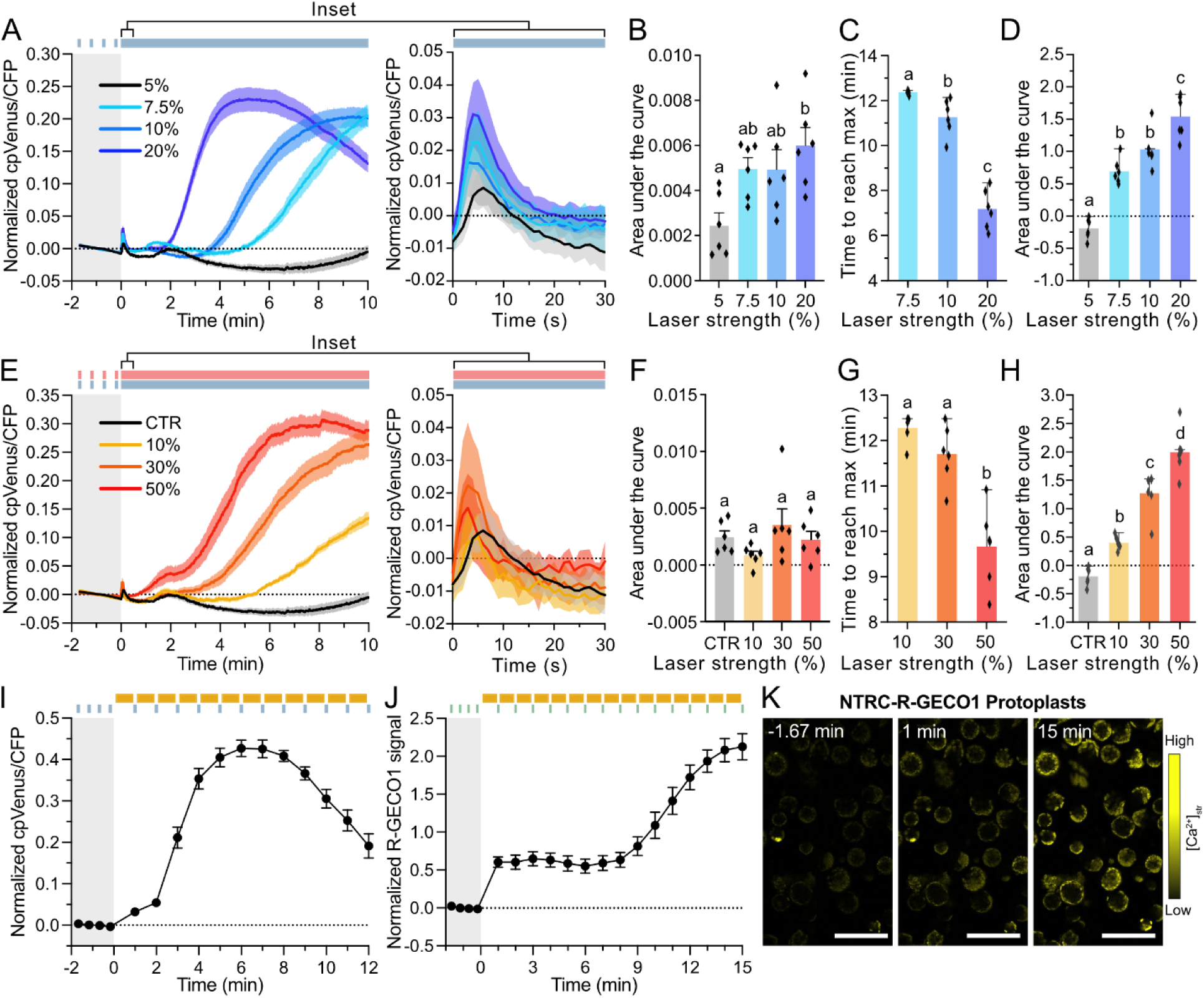
Different light intensities and qualities induce different [Ca^2+^]_str_ signals. **(A)** [Ca^2+^]_str_ dynamics induced by 5% (56 µW), 7.5% (86 µW), 10% (113 µW) and 20% (235 µW) 405 nm laser (blue bar) in cotyledons of the 2BAM4-YC3.6 expressing line. n = 6. The fast [Ca^2+^]_str_ spike in the initial 30 s of HL stimulus is shown in the inset. (**E**) [Ca^2+^]_str_ dynamics induced by 5% (56 µW) 405 nm laser with additional 0 (CTR), 10% (59 µW), 30% (180 µW) and 50% (310 µW) 633 nm laser (red bar) in cotyledons of the 2BAM4-YC3.6 line. CTR and 5% 405 nm laser in (A) are the same data used for comparison. n = 6. (**B, F**) Areas under the curves of the fast [Ca^2+^]_str_ spike in the initial 30 s of HL stimulus. n = 6. (**C, G**) Time to reach maximum values of the persistent [Ca^2+^]_str_ increase in the 10 min of HL stimulus. n = 6. (**D, H**) Areas under the curves of the persistent [Ca^2+^]_str_ increase in the 10 min of HL stimulus. n = 6. (**I**) [Ca^2+^]_str_ dynamics induced by white light (yellow bars) in 2BAM4-YC3.6 line. Cotyledons were intermittently imaged (blue bars) between the white light stimuli. n = 6. (**J**) [Ca^2+^]_str_ dynamics induced by white light in protoplasts of NTRC-R-GECO1 line. n = 11. (**K**) R-GECO1 signals of protoplasts of NTRC-R-GECO1 line, at the time points of 1.67 min before, 1 min after or 15 min after white light stimuli. Different letters above the bars indicate significant differences (1-way ANOVA, p ≤ 0.05). Scale bar, 100 μm. Curves represent mean ± SEM.

**Fig. 4.**
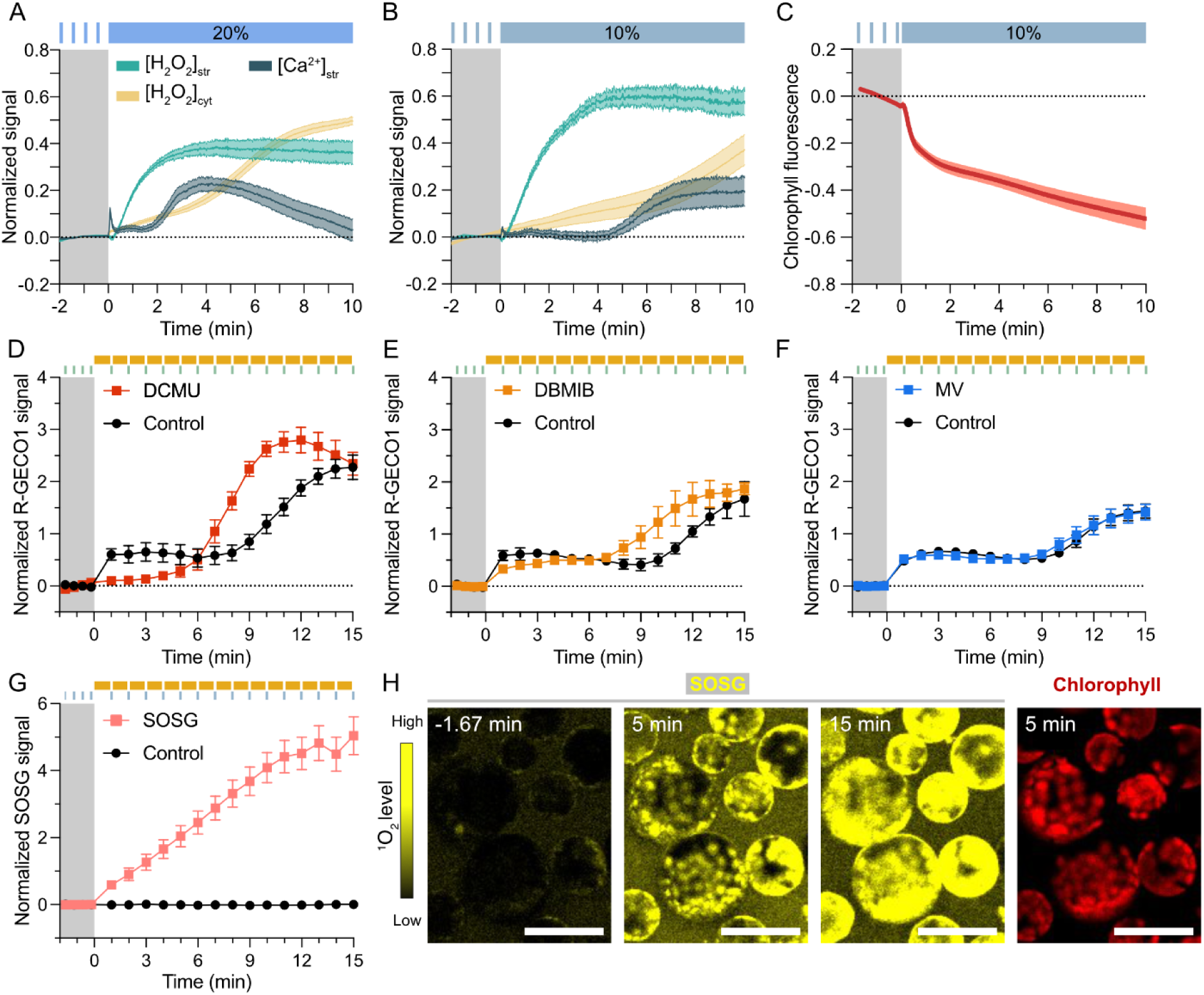
Interaction of [Ca^2+^]_str_ with ROS and photosynthesis. **(A)** [H_2_O_2_]_str_, [H_2_O_2_]_cyt_ and [Ca^2+^]_str_ induced by 20% 405 nm laser light (blue bar) in cotyledons of TKTP-roGFP2-Orp1, cyto-roGFP2-Orp1 and 2BAM4-YC3.6 lines. n = 5. **(B)** [H_2_O_2_]_str_, [H_2_O_2_]_cyt_ and [Ca^2+^]_str_ induced by 10% 405 nm laser (blue bar). n = 5. **(C)** Chlorophyll fluorescence intensity under 10% 405 nm laser (blue bar). n = 7. **(D)** [Ca^2+^]_str_ dynamics induced by white light (yellow bars) in protoplasts of NTRC-R-GECO1 expressing plants. Protoplasts were pre-treated with 10 µM DCMU (n = 5), or 0.1% DMSO (Control, n = 6). The amplitudes of [Ca^2+^]_str_ are shown in the inset. **(E)** [Ca^2+^]_str_ dynamics of NTRC-R-GECO1 line. Protoplasts were pre-treated with 10 µM DBMIB (n = 7), or 0.1% ethanol (Control, n = 3). **(F)** [Ca^2+^]_str_ dynamics of NTRC-R-GECO1. Protoplasts were pre-treated with 50 µM MV (n = 5), or Control (n = 5). **(G)** [^1^O_2_] induced by white light in protoplasts of Col-0. Protoplasts were pre-treated with 100 µM ^1^O_2_ dye SOSG (n = 15), or 1% methanol (Control, n = 3). (**H**) SOSG signal and chlorophyll fluorescence in protoplasts of Col-0. Scale bar, 50 μm. Lines represent mean ± SEM.

Taken together, these data show that the blue light-induced [Ca^2+^]_str_ response originates from the YC3.6 or R-GECO1 reporters and is specific to chloroplasts from cotyledon, first true leaves, as well as mature leaves.

### The stromal Ca^2+^ response is proportional to the light intensity

To determine whether light intensity alters the [Ca^2+^]_str_ response, we varied the strength of the 405 nm laser applied to the 2BAM4-YC3.6 reporter line. Laser power was measured at the focal point on the microscope for the different intensities and summarized in Suppl Table 1. With 5% laser power, an initial spike of [Ca^2+^]_str_ was observed, followed by a small increase lasting from min 3 to 4 (Fig. 3A). However, the later persistent increase of [Ca^2+^]_str_ was absent. Increasing the intensity of blue light to 7.5%, 10%, and 20%, triggered stronger initial [Ca^2+^]_str_spikes (Fig. 3B), and now also the persistent [Ca^2+^]_str_elevations were observed. While the amplitudes of the persistent [Ca^2+^]_str_ elevations remained similar (Fig. 3A), the area under the curve (AUC) increased with the light intensity, implicating a larger [Ca^2+^]_str_accumulation upon higher light intensities during this period of time (Fig. 3C). The time to reach maximum (TRM) of the persistent [Ca^2+^]_str_ increase was significantly shortened with increased light intensity: 20% blue light had the shortest TRM (7.18 min ± 0.36 SEM), followed by 10% blue light (11.26 min ± 0.33 SEM) and 7.5 % blue light (12.37 min ± 0.04 SEM) (Fig. 3D). The TRM of 5% blue light was negative, as there was a heterogeneity of the maximum values appearing in either the first few min or the last few min. Light intensities were in the range, or exceeding, what is generally considered to be HL, e.g. > 1000 µmol photons m^−2^ s^−1^ (Suppl. Table 1). Taken together, the [Ca^2+^]_str_ response required HL intensities and the kinetics correlated with the laser strength.

### Red and white light trigger similar stromal Ca^2+^ responses as blue light does

Next we investigated whether the Ca^2+^ response was specific to blue light or if other wavelengths could also trigger a [Ca^2+^]_str_ increase. In combination with a 5% 405 nm laser, which was necessary for imaging the 2BAM4-YC3.6 reporter but insufficient to trigger persistent [Ca^2+^]_str_ elevations, red light of 633 nm wavelength induced a similar Ca^2+^ responses as blue light (Fig. 3E). The AUC for the fast [Ca^2+^]_str_ spike was not altered by different red light intensities (Fig. 3F), although the amplitude slightly increased, similar to the blue light response. For the persistent [Ca^2+^]_str_ increase, higher 633 nm laser strength led to a shorter TRM (Fig. 3G) and a larger AUC (Fig. 3H). Red light required higher laser strengths for an equal amount of power (watts) as blue light to elicit a similar persistent [Ca^2+^]_str_ increase (Fig. 3, Suppl. Table 1).

To mimic more natural light conditions, we used again white light, a mixture of different light wavelengths across the visible spectrum. In contrast to the transmitted light used previously (Fig. 1D), we now observed a persistent [Ca^2+^]_str_ increase using a mercury short-arc lamp (HXP 120 V) and taking snapshots (∼1 s) of the 2BAM4-YC3.6 signal every 30 s (Fig. 3I), without the need for focusing the light through a glass bead. Unfortunately, the initial [Ca^2+^]_str_ spike occurred too fast and random to observe it with these imaging settings. A halogen light source triggered a persistent [Ca^2+^]_str_ increase as well (Suppl. Fig 5A-C). Similar to what was observed for blue light (Suppl. Fig. 1B), the duration of white light exposure determined the [Ca^2+^]_str_ dynamic. The amplitude was higher after 5 min of light exposure compared to 1.5 min and remained constant thereafter, whereas continuous white light led to a subsequent decrease of [Ca^2+^]_str_ (Suppl. Fig 5A). Under constant white light, the elevated [Ca^2+^]_str_ started to drop from 9 min to 12 min, behaving similar to the [Ca^2+^]_str_ dynamic of 20% blue light (Fig. 3A and 3E). Testing the response on different microscope setups, it became apparent that not all white light intensities triggered a [Ca^2+^]_str_ response. For example, making use of a newly established on-stage illumination system that delivers white LED light to a slide on a microscope [52], no [Ca^2+^]_str_ response could be observed in cotyledons from 2BAM4-YC3.6 expressing plants when switched from 40 µmol photons m^−2^ s^−1^ white light to 600 or 1100 photons µmol m^−2^ s^−1^ (Suppl. Fig. 6). While a transfer from 40 to 1100 µmol photons m^−2^ s^−1^ could be considered HL stress, it appeared not sufficient to trigger a [Ca^2+^]_str_ response. Altogether, our results indicate that a similar perception mechanism exists for HL intensities of either blue, red, or white light, triggering highly similar [Ca^2+^]_str_ dynamics, but a certain threshold of HL intensity is required to trigger the response.

### The high light-induced stromal Ca^2+^ response coincides with ROS production

Since a high intensity of different qualities of light, including white light, triggered the stromal Ca^2^ response, we reasoned that the light stimulus could be perceived through the photosynthesis machinery. Furthermore, the response is triggered in chloroplasts, and not, for example, in root plastids (Fig. 2). Typically, HL exposure generates excessive amounts of photosynthetically derived ROS [17]. So, we visualized ROS accumulation in the cytosol and in the chloroplast stroma with the genetically encoded H_2_O_2_/glutathione sensor roGFP2-Orp1 [45]. While roGFP2 is sensitive to the redox status of the glutathione pool that can be oxidized due to the accumulation of different ROS species (e.g. superoxide, H_2_O_2_), the roGFP2-Orp1 fusion protein has specificity for H_2_O_2_ as an oxidant and the glutathione system as reductant, *in vivo* [46, 57]. We imaged the 2BAM4-YC3.6 and roGFP2-Orp1 reporters with the same microscopy settings in order to compare the accumulation of ROS and Ca^2+^ over time. Indeed, the application of 20 % blue light triggered pronounced H_2_O_2_ accumulations in the cytosol and chloroplast stroma (Fig. 4A). The initial [Ca^2+^]_str_ spike preceded stromal H_2_O_2_ accumulation, whereas the persistent [Ca^2+^]_str_ increase occurred almost 2 min following H_2_O_2_ accumulation. H_2_O_2_ accumulated in the cytosol as well, albeit at a slower pace than in the chloroplast stroma, as it was previously reported for white HL [17]. Notably, 10 % blue light triggered a clear stromal H_2_O_2_ accumulation but a much weaker [Ca^2+^]_str_ increase (Fig. 4B). Similar responses were recorded with the genetically encoded glutathione redox potential sensor, GRX1-roGFP2 [45] (Suppl. Fig. 7).

It became apparent over the course of our experiments that the light regime applied to cotyledons was quenching the chlorophyll autofluorescence (Fig. 4C), which is potentially indicative of enhanced non-photochemical quenching [9] or damage to the photosystems [14]. To confirm the involvement of the photosynthetic electron transport reactions in the [Ca^2+^]_str_ response, we tested the effect of 3-(3,4-dichlorophenyl)-1,1-dimethylurea (DCMU), that irreversibly blocks the Q_B_ plastoquinone binding site of photosystem II, obstructing the electron flow from photosystem II to plastoquinone. DCMU experiments were performed using protoplasts instead of leaves, as penetrance of chemicals is more effective in protoplasts, as shown for example with the mitochondrial dye (Suppl. Fig. 2D). In DCMU-treated protoplasts from the NTRC-R-GECO1 line, white light-induced persistent [Ca^2+^]_str_ increase reached a maximum earlier than control (Fig. 4D). However, the amplitude of the [Ca^2+^]_st_ response remained unaffected. Interestingly, the pH-dependent increase in R-GECO1 signal before the persistent [Ca^2+^]_str_ increase was abolished, indicating that as expected [17], light-induced stromal alkalinization was inhibited by DCMU treatment. Dibromothymoquinone (DBMIB), a plastoquinone analogue and alternative inhibitor that blocks binding at the Q_o_-site of the Cyt b_6_f complex, also led to an earlier [Ca^2+^]_str_ accumulation, albeit to a lesser extent than DCMU (Fig. 4E). Treatment with methyl viologen (MV), which leads to superoxide (O_2_•–) production by funneling electrons from photosystem I to oxygen (O_2_), did not markedly alter the [Ca^2+^]_str_ response observed in untreated protoplasts (Fig. 4F).

As there was a significant delay between stromal H_2_O_2_ accumulation and [Ca^2+^]_str_ increase, especially at lower light intensity (Fig. 4B), and DCMU treatment accelerated the stromal Ca^2+^ response, we tested the potential involvement of other photosynthetically derived ROS. DCMU prevents the reduction of plastoquinone at photosystem II, acting upstream of the inhibitory effect of DBMIB at the site of plastoquinone oxidation by the cytochrome b_6_f complex [58]. Hence, if ROS were involved in the HL-induced [Ca^2+^]_str_ response, they would likely be generated early during the photosynthetic electron transport, such as ^1^O_2_ at photosystem II [59]. Indeed, HL triggered a consistent increase of ^1^O_2_ in chloroplasts from NTRC-R-GECO1 protoplasts visualized with the chemical Singlet Oxygen Sensor Green (SOSGreen) (Fig. 4G and 4H). SOSGreen is highly specific for ^1^O_2_, but has also a tendency for photosensitization in white light leading to an increased background signal [60]. This effect was visible in our experiment. Hence, background subtraction was performed for quantification. Furthermore, the signal that overlapped with chloroplast localization was stronger than the background. In conclusion, the observed [Ca^2+^]_str_ response is likely the result of a HL reaction at the photosynthetic complexes, and might interact with photosynthetically derived ROS, especially ^1^O_2_.

### Stromal Ca^2+^ elevations are caused by light, rather than by the temperature component of HL

Previously, heat shock was found to induce [Ca^2+^]_str_ elevations [61]. To distinguish whether the HL-induced [Ca^2+^]_str_ elevations are the result of light absorption or a potential temperature effect, we conducted HL experiments under different temperature regimes. As observed previously with a luminescent aequorin reporter by Lenzoni & Knight [61], at 40℃, the initial [Ca^2+^]_str_ level was elevated likely because of heat shock (Fig. 5A). However, [Ca^2+^]_str_ dropped upon continuous HL treatment and increased afterwards (Fig. 5A). Non-normalized data illustrate the difference between the initial [Ca^2+^]_str_ at 40℃ with those at lower temperatures (Fig. 5A). At 10℃, a delayed [Ca^2+^]_str_ increase was observed (Fig. 5A). So, temperature did interfere with the HL-induced persistent [Ca^2+^]_str_ elevations, as [Ca^2+^]_str_ rose faster along with temperature increase (Fig. 5A), albeit not to the extent observed at 40℃. Moreover, the [Ca^2+^]_str_ dynamics looked overall similar to those observed previously without temperature control.

**Fig. 5.**
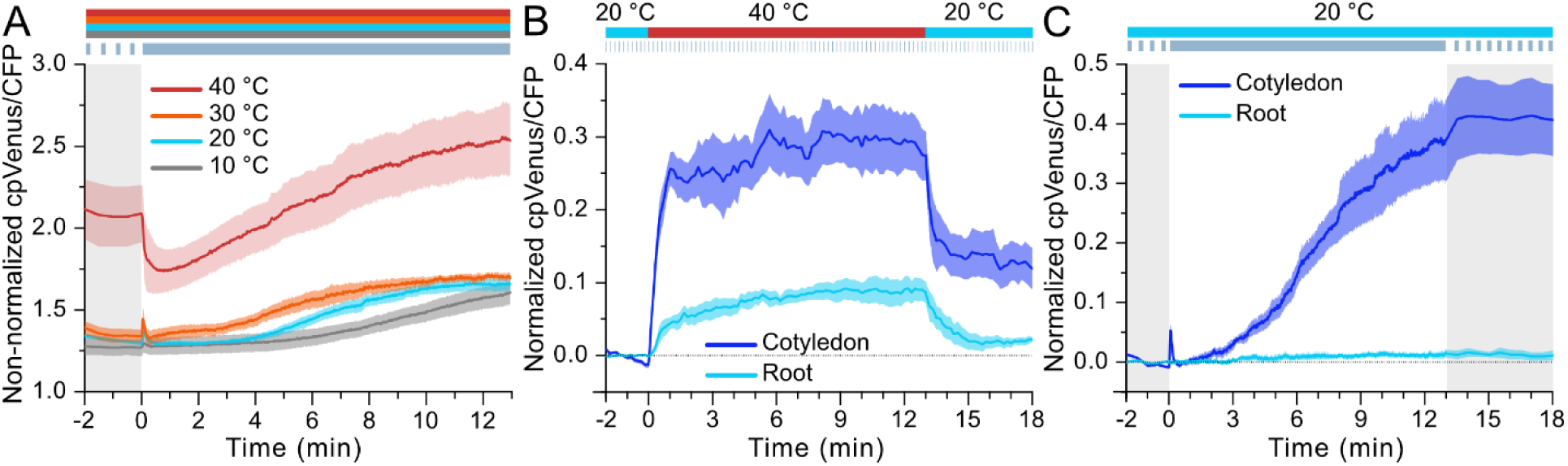
Interaction of HL-induced [Ca^2+^]_str_ with temperature. **(A)** [Ca^2+^]_str_ dynamics induced by 10% 405 nm laser (dark blue bar) in cotyledons of the 2BAM4-YC3.6 expressing plants at 10℃ (grey bar), 20 ℃ (light blue bar), 30℃ (orange bar) and 40℃ (red bar). n = 5. (**B**) [Ca^2+^]_str_ dynamics of 2BAM4-YC3.6 expressing cotyledons/roots under a temperature shift (20℃ to 40℃ to 20 ℃). n = 5. (**C**) [Ca^2+^]_str_ dynamics of 2BAM4-YC3.6 cotyledons/roots under light shift (intermittent imaging to continuous imaging to intermittent imaging). n = 5. The temperature regime was controlled by a CherryTemp heating/cooling system. Lines represent mean ± SEM.

To further distinguish the light effect from a potential temperature effect of HL, we performed simultaneous temperature and light shift experiments. We observed an increase of [Ca^2+^]_str_upon heat shock (40℃) in root cells (Fig. 5B), while a light intensity that triggers [Ca^2+^]_str_in green tissues had no effect on [Ca^2+^]_str_ in roots (Fig. 5C). This indicates that our experimental HL system does not cause a strong temperature effect, at least not in roots. Furthermore, for cotyledons, the Ca^2+^ signatures triggered by heat shock and HL were very different: the initial [Ca^2+^]_str_ spike was absent following heat shock and [Ca^2+^]_str_ rose more slowly during HL compared to heat shock-induced [Ca^2+^]_str_ elevations (Fig. 5B and 5C). Importantly, [Ca^2+^]_str_ remained high after switching from HL to darkness under continuous cooling at 20℃ (Fig. 5C), while the [Ca^2+^]_str_ levels dropped when switching to 20℃ after heat shock (Fig. 5B). The same imaging setting of the temperature shift experiment was carried out under constant 20℃, ensuring that the imaging setting itself did not trigger any [Ca^2+^]_str_changes (Suppl. Fig. 8). In conclusion, temperature influenced the speed at which the persistent [Ca^2+^]_str_ took place. However, temperature only mildly affected the light response, unlike the drastic response to heat shock at 40℃.

### The high light-induced stromal Ca^2+^ response likely originates from the ER rather than from the cytosol

Calcium signaling relies on the storage of Ca^2+^ in subcellular compartments such as the apoplast, vacuole and the endoplasmic reticulum (ER), and its release to cellular localities with low Ca^2+^ concentration. Subsequent active transport of Ca^2+^ back to the stores leads to typical peaks of free [Ca^2+^] - so-called calcium signatures [21, 62]. To investigate the origin(s) of the HL-induced stromal Ca^2+^, we subjected a cytosolic Ca^2+^ sensor line, NES-YC3.6 [43], to HL treatment. Blue light triggered [Ca^2+^]_cyt_ peaks at laser intensities that were previously applied to the 2BAM4-YC3.6 reporter line (Fig. 6A). However, contrary to the [Ca^2+^]_str_ response, a clear [Ca^2+^]_cyt_ response was triggered already at 5% laser strength (Fig. 6B and 6C). Besides, the [Ca^2+^]_cyt_ peak was slower than the fast [Ca^2+^]_str_ spike and preceded the later persistent [Ca^2+^]_str_ increase by several minutes (Fig. 6D). Red light triggered a persistent [Ca^2+^]_str_ increase, but did not trigger a pronounced [Ca^2+^]_cyt_ peak (Fig. 6E). White light triggered a clear [Ca^2+^]_cyt_ peak as well (Suppl. Fig. 5B). We crossed the NTRC-R-GECO1 line with a cyto-nuclear targeted GCaMP3 line [47] and subjected it to HL, confirming the observations made with single chloroplast and cytosol-targeted YC3.6 lines (Suppl. Fig. 10). Assuming the HL-induced [Ca^2+^]_str_ response is not directly linked to a [Ca^2+^]_cyt_ peak, alternative sources for the [Ca^2+^]_str_ must be considered.

**Fig. 6.**
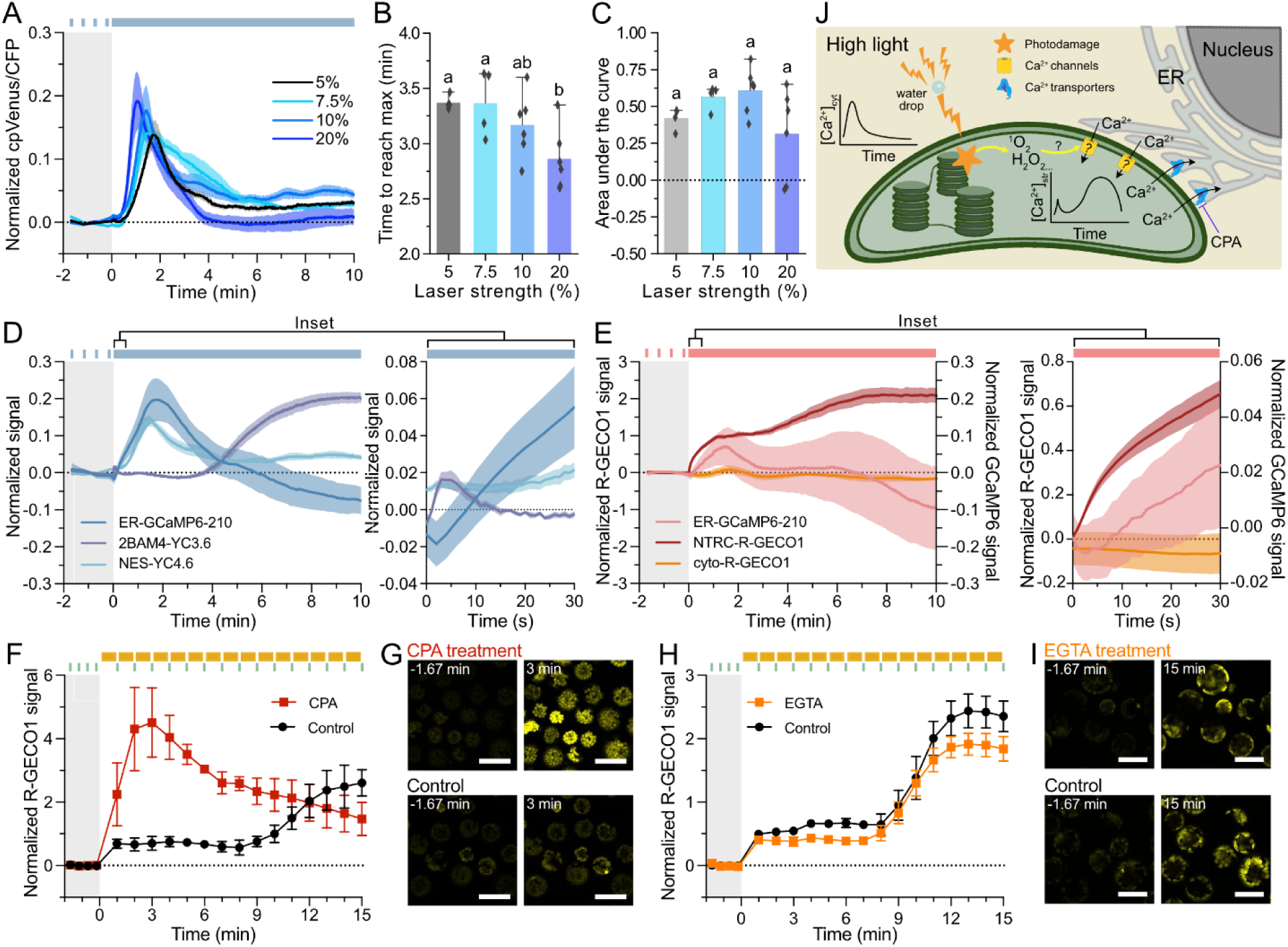
[Ca^2+^] dynamics in other compartments. **(A)** [Ca^2+^]_cyt_ dynamics induced by 5% (56 µW, n = 4), 7.5% (86 µW, n = 4), 10% (113 µW, n = 6) and 20% (235 µW, n = 6) 405 nm laser (blue bar) in cotyledons of NES-YC3.6 expressing plants. (**B**) Time to reach maximum values of [Ca^2+^]_cyt_ in the 10 min of HL stimulus. (**C**) Areas under the curves of [Ca^2+^]_cyt_ in the 10 min of HL stimulus. (**D**) [Ca^2+^]_str_ and [Ca^2+^]_cyt_ induced by 10% 405 nm laser (blue bar) in cotyledons of 2BAM4-YC3.6 (n = 6) and NES-YC3.6 lines (n = 5). [Ca^2+^]_ER_ induced by sequential 10% 405 nm / 20% 488 nm laser (blue bar) in cotyledons of ER-GCaMP6-210 line (n = 6). [Ca^2+^] in the initial 30 s of HL stimulus was shown in the inset. (**E**) [Ca^2+^]_ER_, [Ca^2+^]_str_ and [Ca^2+^]_cyt_ induced by 30% 633 nm laser (red bar) in cotyledons of ER-GCaMP6-210 (n = 8), NTRC-R-GECO1 (n = 5) and cyto-R-GECO1 expressing lines (n = 5). (**F**) [Ca^2+^]_str_ dynamics induced by white light in protoplasts of NTRC-R-GECO1 line. Protoplasts were pre-treated with 25 μM CPA (n = 4), or 0.25% DMSO (Control, n = 5). (**G**) R-GECO1 signals of protoplasts of NTRC-R-GECO1 line under 25 μM CPA treatment or Control, at the time points of 1.67 min before and 3 min after white light stimuli. (**H**) [Ca^2+^]_str_ dynamics of NTRC-R-GECO1 line. Protoplasts were pre-treated with 10 mM EGTA (n = 5), or control (n = 3). (**I**) R-GECO1 signals of protoplasts of NTRC-R-GECO1 line under 10 mM EGTA treatment or Control, at the time points of 1.67 min before and 15 min after white light stimuli. (**J**) Hypothetical model of ER-to-chloroplast-calcium transfer. Different letters above the bars in (B) and (C) indicate significant differences (p ≤ 0.05). Scale bar, 50 μm. Lines represent mean ± SEM.

The ER is one of the major Ca^2+^ stores in plant cells, and close contacts between the ER and chloroplasts have been previously observed [63–65]. To test the potential involvement of ER in the HL-induced [Ca^2+^]_str_ response, we subjected the ER Ca^2+^ sensor line ER-GCaMP6-210 [66] to blue light exposure under similar conditions as NES-YC3.6 and 2BAM-YC3.6 (Fig. 6D). In contrast to the cytosol, the [Ca^2+^]_ER_ response mirrored the stromal response: [Ca^2+^]_ER_ dropped at the same time as the initial [Ca^2+^]_str_ peaked (Fig. 6D inset), after which it slowly recovered, to decrease again during the persistent [Ca^2+^]_str_ increase. Simultaneously, the blue light-induced [Ca^2+^]_cyt_ peak was closely followed by a rise of the [Ca^2+^]_ER_, which could indicate a transfer of cytosolic Ca^2+^ to the ER lumen during blue light exposure. The GCaMP6 family of fluorophores display dual excitation peaks at 410 nm and 474 nm, corresponding to Ca^2+^-independent and Ca^2+^-dependent emissions, respectively [49, 50]. This property enabled ratiometric imaging by sequential excitation at 405 and 488 nm (Suppl. Fig. 11A), excluding the impact of fluorophore photobleaching (Suppl. Fig. 11B). The ER-GCaMP6-210 signal showed a similar response to red light exposure, but the decrease was less pronounced compared to blue light (Fig. 6E). This could be the consequence of an overall weaker [Ca^2+^]_str_ response to 30% red light compared to 10% blue light (Fig. 3A and 3E). Further evidence for a potential Ca^2+^ transfer from ER to chloroplast stroma during HL exposure came from chemical treatments with cyclopiazonic acid (CPA), an inhibitor of type IIA ER-type Ca^2+^-ATPases (ECAs) [67] that was shown to deplete [Ca^2+^]_ER_ [66]. When subjecting protoplasts to a white HL stimulus, CPA treatment greatly enhanced the initial [Ca^2+^]_str_ spike and abolished the persistent [Ca^2+^]_str_ increase compared to a DMSO control (Fig. 6F and 6G). Chelation of extracellular Ca^2+^ with EGTA had little effect on the persistent [Ca^2+^]_str_ increase, ruling out a direct Ca^2+^ transfer from the extracellular space to the chloroplast stroma (Fig. 6H and 6I). A decrease in amplitude of the persistent [Ca^2+^]_str_ increase is consistent with the observations by Huang et al. [68], who observed that extracellular Ca^2+^ chelation with EGTA leads to a 20% reduction in release of [Ca^2+^]_ER_ to the cytosol. Although we cannot rule out the contribution from other Ca^2+^ stores such as the vacuole and the Golgi apparatus, considering the distinct [Ca^2+^] dynamics in chloroplast, cytosol, and ER, together with the strong effect of a known [Ca^2+^]_ER_ inhibitor, it is likely that Ca^2+^ is transferred directly from ER to the chloroplast stroma during HL exposure.

## Discussion

HL stress is a major obstacle for plants to achieve maximal photosynthetic efficiency and thereby optimal growth. Thus, it is important for plants to sense, respond and acclimate to HL stress [8]. Our study reveals a novel Ca^2+^ response pathway to sense and respond to excessive light exposure that eventually could lead to HL stress acclimation. A biphasic [Ca^2+^]_str_ increase was triggered in green tissues by different light regimes and qualities, including blue, red, and white light, either focused through a glass bead or not. Increasing light intensity and temperature led to an accelerated response, albeit with a different dynamic than heat shock. The [Ca^2+^]_str_ increase was accompanied by a decrease in chlorophyll autofluorescence and accumulation of ROS, clear hallmarks of photodamage to the electron transport chain. Finally, although HL induced an increase in nucleocytoplasmic [Ca^2+^], direct measurements of [Ca^2+^]_ER_ levels and inhibitor treatments suggest that the HL-induced [Ca^2+^]_str_ rise is primarily due to Ca^2+^ transfer between the ER and chloroplast.

### The fast initial [Ca^2+^]_str_ spike precedes all other Ca^2+^ signals and ROS accumulation

In both the 2BAM4-YC3.6 line and NTRC-R-GECO1 lines, HL induced a fast initial [Ca^2+^]_str_ spike that lasted about 30 sec in 2BAM4-YC3.6 line and 2 min in the NTRC-R-GECO1 line. The slow decay of the initial [Ca^2+^]_str_ spike in the NTRC-R-GECO1 line is a consequence of the fast alkalinization upon illumination, indicated by the chloroplast-targeted pH sensor cp-cpYFP (Suppl. Fig. 3A). Alkalinization was reported to promote R-GECO1 fluorescence [44], and here, masked the fast initial [Ca^2+^]_str_ spike in NTRC-R-GECO1 because of averaging the signal intensities from all chloroplasts. However, analysis of individual chloroplasts in the images revealed a spike that originated from a few random chloroplasts (Fig. 2F; Suppl. Fig. 3C, 3D), similar to what we saw for the 2BAM4-YC3.6 line. It is unlikely that ROS are involved, since the initial [Ca^2+^]_str_ spike precedes the accumulation of H_2_O_2_ (Fig. 4A and 4B) and ^1^O_2_ (Fig. 4G). Furthermore, the initial [Ca^2+^]_str_ spike was independent of light quality (Fig. 3A and E) and somewhat of light intensity (Fig. 3B and 2F). At least, a different mechanism is likely responsible for the later persistent [Ca^2+^]_str_ increase, which required a higher light intensity than 5% blue laser strength and depended largely on light intensity (Fig. 3), and for the [Ca^2+^]_cyt_ peak, which is triggered mainly by blue light (Fig. 6). One potential explanation stems from the ER and CPA experiments: The initial [Ca^2+^]_str_ spike is mirrored by a dip in [Ca^2+^]_ER_ (Fig. 6D) and CPA treatment drastically enhanced the initial [Ca^2+^]_str_ spike (Fig. 6F). Hence, the initial [Ca^2+^]_str_ spike could be the product of an imbalance between a known light-induced stromal Ca^2+^ influx [75–77] and a hypothetical Ca^2+^ efflux to the ER, mediated by a downregulation of ER-resident Ca^2+^ ATPase activity. Further research is required to understand the mechanistic details of why these chloroplasts are triggered and not others, and to test a potential physiological impact of this signal.

### The persistent [Ca^2+^]_str_ increase is likely caused by severe HL stress, accompanied by photoinhibition and ^1^O_2_ accumulation

A slower but persistent [Ca^2+^]_str_ increase followed the fast initial [Ca^2+^]_str_ spike in both the 2BAM4-YC3.6- and NTRC-R-GECO1-expressing lines. Although R-GECO1 is pH sensitive, the pH drops during the later stage of illumination (Suppl. Fig. 3A), indicating that the slow increase of NTRC-R-GECO1 signal was due to an authentic increase of Ca^2+^, validating the 2BAM4-YC3.6 result. The persistent [Ca^2+^]_str_ increase has all the hallmarks of a HL response. Although we have not measured photosynthetic activity directly (e.g. Fv/Fm) due to technical constraints of the small imaging surface that is probed via microscopy, the clear decrease of chlorophyll autofluorescence (Fig. 4C) hints at a strong impairment of the photosystems, where PSII is the main target of photoinhibition [15]. Furthermore, there is a clear dose response of light to trigger the persistent [Ca^2+^]_str_ increase (Fig. 3) that occurs only when the light intensity exceeds 1100 µmol photons m^−2^ s^−1^ (Suppl. Fig. 6). The threshold to trigger a robust [Ca^2+^]_str_ response is around 2000 µmol photons m^−2^ s^−1^ (Fig. 3, Suppl. Table 1), which is the light intensity typically found outdoors under a bright sunny sky. Going beyond exposure to light of 2000 µmol photons m^−2^ s^−1^, natural lensing conditions should be considered, for example through water droplets. The effect of natural light focusing on plant physiology is surprisingly little described [5], however, spots of focused sunlight are a rather common sight in nature, for example on days with intermittent rain and sunshine (Fig. 1A and B). We have mimicked natural light lensing by focusing the transmission light of the microscope through a glass bead to elicit a [Ca^2+^]_str_ increase (Fig. 1C). Specialized acclimation signaling, for example through a burst of stromal [Ca^2+^], could be required to avoid sunburn and potential (runaway) cell death [80, 81] and should be the focus of further study.

Intriguingly, a brief pulse of HL caused a sustained [Ca^2+^]_str_ increase that remained elevated, unlike the response to continuous HL over a comparable time period (Suppl. Fig. 1B and Suppl. Fig. 5B). The cause for this difference is unclear, however, it is in line with the previous observation that a pulse of 2-5 min HL leads to a significantly higher level of HL-acclimation related gene expression than 10 min of HL exposure [82].

Contrary to previous research suggesting a potential role of HL-induced H_2_O_2_ production in the [Ca^2+^]_str_ response [41, 42], our findings show that the stromal H_2_O_2_ dynamics, measured using roGFP2-Orp1-expressing lines, are not correlated with the [Ca^2+^]_str_ response in terms of both kinetics and light intensity requirements (Fig. 4A and 4B). One counterargument is that a certain threshold of H_2_O_2_ flux may be required to induce the persistent [Ca^2+^]_str_ increase and that the sensor relies on endogenous E_GSH_ for re-generation, which is itself variable due to its various functions. Additionally, it was previously shown that exogenous H_2_O_2_ application leads to a prolonged [Ca^2+^]_str_ elevation [24, 41]. Furthermore, it is important to note that roGFP2-Orp1 detects both H_2_O_2_ and lipid hydroperoxides (LOOH) [83], which may complicate the interpretation of our data. Recently, Exposito-Rodriguez et al. [83]. demonstrated that a mutated version of the sensor, roGFP2-synORP1, is less affected by LOOHs, and that HL treatment results in similar accumulations of H_2_O_2_ and LOOHs in the chloroplast stroma [83]. Notably, during severe photoinhibition (a combination of HL and DCMU treatment), there was a greater LOOH accumulation in the cytosol.

Although the involvement of H_2_O_2_ cannot be entirely ruled out, chemical manipulation of photosynthetic electron flux downstream of photosystem II supports the involvement of other ROS such as ^1^O_2_. Indeed, DCMU and DBMIB speed up the persistent [Ca^2+^]_str_ increase (Fig. 4D and 4E), while previously these treatments were shown to decrease H_2_O_2_ accumulation [17, 83]. Furthermore, MV treatment was previously shown to increase roGFP2-Orp1 signal more strongly in the chloroplast than in the cytosol [45], however in our study MV did not alter the HL-induced [Ca^2+^]_str_ increase (Fig. 4F). A combination of HL and DCMU leads to increased ^1^O_2_ production at photosystem II and the resulting production of LOOHs through the oxidation of lipids by ^1^O_2_ [83]. In our study, we observed a steady increase in ^1^O_2_, measured using the SOSG (Singlet Oxygen Sensor Green) chemical probe that coincides more closely with the persistent [Ca^2+^]_str_ elevation (Fig. 4D and 4G).

### Benefits and limitations of the new chloroplast-targeted GECI, NTRC-R-GECO1

R-GECO1 has certain advantages over YC3.6. The yellow-green (561 nm) excitation of R-GECO1 should affect the photosynthetic light reactions less than blue light as long as the yellow-green measurement light is softer than the blue light used to trigger the [Ca^2+^]_str_ increase [69]. Furthermore, 561 nm excitation light does not substantially overlap with the absorption spectra of UV-B, blue and far-red plant photoreceptors (i.e, UVR-8, cryptochromes, phototropins, and the Pfr form of phytochrome), and only partially with the Pr form of phytochrome [70]. The selected NTRC-R-GECO1 reporter line is not sensitive to silencing (at least to the fourth generation) and expressed uniformly in wild type Col-0 background. In contrast, 2BAM4-YC3.6 is expressed in an *rdr6* background that is deficient for RNA-DEPENDENT RNA POLYMERASE 6, important for siRNA biogenesis [71], as it is prone to silencing [43]. While this is not a problem per se, *rdr6* has a downwards curled leaf phenotype [72], and the additional mutant allele complicates the use of the 2BAM4-YC3.6 reporter in mutant backgrounds either by crossing or genetic transformation. R-GECO1 lacks ratiometric capacity, however, has a large dynamic range, making the [Ca^2+^]_str_ response highly visible (Suppl. Movie 2, 3, and 4) and a single fluorophore simplifies and democraties the visualization (e.g. on low-end microscopes or stereoscopes) and use in combination with other reporters (e.g. GCAMP3, Suppl. Fig. 9A-C) [73]. Other disadvantages are that R-GECO1 has an undesirable blue-light-activated photo switching behavior [74] and is more pH-sensitive than YC3.6 [44]. Both effects were likely responsible for the initial rise in R-GECO1 fluorescence at the start of the continuous blue light trigger (Fig. 2A). However, after the initial rise, a [Ca^2+^]_str_ increase was observed similar to the secondary transient increase for 2BAM4-YC3.6 (Fig. 2A). Furthermore, the pH effect is negligible compared to the signal increase of R-GECO1 during a Ca^2+^ spike (Suppl. Fig 3D).

### HL-induced [Ca^2+^]_str_ kinetics differ between the epidermis, stomata and the mesophyll

For both 2BAM4-YC3.6 and NTRC-R-GECO1 expressing plants, the persistent [Ca^2+^]_str_ increase appeared later in mesophyll and stomata compared to the epidermis (Fig. 2B and 2E). Although the reason for this is unclear at the moment, it could be connected to a recently recognized class of plastids, specialized for stress sensing: sensory plastids [84]. Sensory plastids can be found in the epidermis, are smaller, produce more stromules, have a less complex grana stacking and contain fewer plastoglobules than mesophyll chloroplasts. They are distinguished from mesophyll chloroplasts by a different proteome with stress associated features. Interestingly, Beltran et al. reported a change in gene expression for calcium-mediated signaling pathways, including numerous calmodulin and rapid alkalinization factor-like signal elements in cells containing sensory plastids [85]. The presence of these sensory plastids in the epidermis may explain the distinct HL-induced [Ca^2+^]_str_ kinetics observed compared to mesophyll and stomata, potentially playing a key role in signaling during HL stress in plants. However, at this point we cannot rule out that light penetration differs between tissues/cell types along a depth gradient in the leaf and that the mere position of chloroplasts relative to the leaf surface determines the [Ca^2+^]_str_ response. Light diffusion and perception in plants are further complicated by air pockets in the leaf mesophyll and hypocotyl, which scatter incoming light creating a distinct light gradient within green tissues [86]. Nawkar et al. identified an *abcg5* mutant that makes the hypocotyl more transparent due to water-filled air pockets. This causes a light tropism defect that can be mimicked by infiltrating water into the hypocotyl and leaf [86]. Notably, infiltration of water impaired the persistent [Ca^2+^]_str_ increase, thereby implicating light scatter in the HL response (Suppl. Fig. 10). This effect may be further influenced by heterogeneity of the leaf internal light environment [87], and by the chloroplast avoidance response, in which chloroplasts move from the surface to the side of the cell to shade each other from HL [88]. While light penetration differences may explain the variation between epidermal and mesophyll chloroplasts, it does not account for the delayed response in stomatal chloroplasts based solely on leaf depth.

### The HL-induced [Ca^2+^]_cyt_ spike is an effect of a blue component in the white light spectrum

Apart from the observation of HL-induced [Ca^2+^]_str_ dynamics, we observed a clear blue and white light induced [Ca^2+^]_cyt_ peak (Fig. 6A and Suppl. Fig. 1B, 5D and 5E). Blue light triggered a stronger [Ca^2+^]_cyt_ peak compared to red light (Fig. 6E) and contrary to the [Ca^2+^]_str_ dynamics, TRM and AUC of the [Ca^2+^]_cyt_ peak were similar for all blue light intensities used in this study (Fig. 6B and 6C). This indicates that the potential light sensing mechanism for the [Ca^2+^]_cyt_ peak depends on a blue light receptor and likely it is the blue spectrum in the white light treatments that induce the response. These transient increases of cytosolic Ca^2+^ may come from phototropin (a blue light photoreceptor), which was reported to mediate blue light induced [Ca^2+^]_cyt_ increase [89–91].

Interestingly, Baum et al. [91] reported that blue light exposure does not trigger a [Ca^2+^]_str_ increase. However, their experimental setup precluded the possibility of observing the [Ca^2+^]_str_ dynamics that we report. Baum et al. triggered Ca^2+^ responses with an external blue light source (fluence rate of 600 μmol⋅m−2⋅s−1) for 10 sec, plus 3-5 sec to allow for auto fluorescence decay, after which they measured [Ca^2+^]_str_ with aequorin [91]. Most likely, the initial fast [Ca^2+^]_str_ peak was already triggered during the blue light stimulus and went unobserved as they measured only after the blue light stimulus. Furthermore, the measurements following the light stimulus were either too short to observe the persistent [Ca^2+^]_str_ increase, or the light stimulus was not strong enough (since the [Ca^2+^]_cyt_ peak requires less intense light). Note that the fast [Ca^2+^]_str_ peak is already triggered at 5% blue light, so in theory Baum et al. might have observed it, if they had chloroplast-targeted fluorescent GECIs at the time.

A profound alteration of nuclear gene expression and translation occurs during acclimation to HL stress [82, 92, 93]. The observation of a white HL-induced [Ca^2+^]_cyt_ increase opens the door for the downstream investigation of a direct effect of this Ca^2+^ signal on nuclear gene expression. Recently, Moore et al, identified a translation-dependent retrograde signaling network that is regulated by MPK6, AKIN10, SAP2/3 and CML49 within minutes of HL treatment [92]. MPK6 and AKIN10 were found to control a HL-induced cytosolic translational network followed by the nuclear relocation of a SAP2/3 and CML49 complex and a transcriptional loop of the HL response. MPK6 can be activated through an unknown mechanism in response to Ca^2+^ signals [1] and CML49 contains two predicted functional calcium binding EF-hands that can bind Ca^2+^ directly. Since the GCaMP3 reporter in the NTRC-R-GECO1 crossed line is not excluded from the nucleus (whereas the NES-YC3.6 reporter is), we could observe a [Ca^2+^] spike also in the nucleus following HL (Suppl. Fig. 9A) and in a previously established cyto-nuclear localized R-GECO1 reporter line (Suppl. Fig. 9D and Movie 4). HL-induced systemic [Ca^2+^] elevations were reported previously, but lacked subcellular information due to the use of the Ca^2+^ reporter dye FLUO4-AM [94]. Fichman et al. likely missed observing the local HL-induced [Ca^2+^]_cyt_ peak as the Ca^2+^ response was measured only after two min of HL treatment. Hence, MPK6 and CML49 are potential targets of the HL-induced nucleo-cytosolic [Ca^2+^] spike we have observed or the previously observed systemic Ca^2+^ response [94], leading to altered transcription and translation for HL acclimation.

### The HL-induced persistent [Ca^2+^]_str_ increase is conserved throughout the green lineage

Similar HL-induced [Ca^2+^]_str_ responses were recently observed in the green algae Chlamydomonas and the diatom Phaeodactylum [95, 96]. The HL treatments differ from this study, i.e a HL pulse of 90 s in the study by Pivato et al. [41] and an intermittent excitation every 4 s at 2365-4000 μmol photons m^−2^ s^−1^ by Flori et al. [42]. The impact of HL stress on the functioning of photosynthesis seems to be at the origin of the stromal Ca^2+^elevation in the three organisms studied. Indeed, the use of chemicals inhibiting the transport of electrons within the photosynthetic chain (DCMU in Arabidopsis and Chlamydomonas and MV in Phaeodactylum), thus promoting the production of ROS, was correlated with the increase in Ca^2+^ in the stroma. The H_2_O_2_ measurements carried out under HL stress conditions in the three studies also reinforce this correlation of increased stromal Ca^2+^ when the photosynthetic apparatus is damaged by stress. Neither the initial fast [Ca^2+^] spike nor the cytosolic [Ca^2+^] changes were observed in Chlamydomonas and Phaeodactylum, during HL treatment. The reasons behind could be the different setup and/or methodology. Alternatively, vascular plants like Arabidopsis may have acquired a more complex HL induced Ca^2+^ signature while retaining the persistent stromal response. We suggest that a potential HL acclimation mechanism mediated by Ca^2+^ in the chloroplast has conserved evolutionary roots.

### A potential origin and mechanism of HL-induced [Ca^2+^]_str_ response

To understand how the HL-induced [Ca^2+^]_str_ signature is formed and to eventually find a way to disrupt it, we set out to pinpoint the potential store(s) from where the Ca^2+^ originates. The cytosol acts as a reservoir from where Ca^2+^ peaks are transferred into the chloroplast stroma, exemplified by a short delay between cytosol and chloroplast peaks in response to external elicitors (e.g. salt, mannitol, flg22) [10, 19]. Furthermore, Ca^2+^ is believed to be transported from the cytosol to the stroma under normal (not high) light conditions, although this does not lead to a stromal [Ca^2+^] accumulation, as Ca^2+^ is transported concurrently to the thylakoid [4, 16, 20]. Two points from our study preclude a direct transfer of the HL-induced [Ca^2+^] peak from cytosol to stroma that would result in the persistent [Ca^2+^]_str_ increase: i) The disparity in timing between the [Ca^2+^]_cyt_ peak and persistent [Ca^2+^]_str_ increase (Fig. 6D and Suppl. Fig. 1B), and ii) the absence of a clear red light induced [Ca^2+^]_cyt_ peak at a light intensity that triggers the persistent [Ca^2+^]_str_ increase (Fig. 6E). Furthermore, HL does not trigger a [Ca^2+^]_cyt_ peak in Chlamydomonas and Phaeodactylum although there is a clear [Ca^2+^]_str_ increase [41, 42]. However, we cannot rule out the possibility that Ca²⁺ is transferred to the stroma during the [Ca^2+^]_cyt_ peak without causing a detectable increase in [Ca^2+^]_str_. This could occur if Ca^2+^ is rapidly pumped into the thylakoid or other compartments during the lag time or ‘valley’ between the initial [Ca^2+^]_str_ spike and subsequent persistent increase. In this valley, photosynthesis is still functional, evidenced by stromal alkalinization (Suppl. Fig. 3B), which might drive Ca^2+^ away from the stroma into the thylakoid or other subcellular locations.

We cannot exclude a potential contribution of the thylakoid lumen to HL-induced [Ca^2+^]_str_ dynamics. Solid evidence was presented for the light-dependent uptake of Ca^2+^ into the thylakoid lumen in exchange for H^+^ [97], likely mediated by BICAT1 [29]. Ettinger et al. have speculated that Ca^2+^ is released from the thylakoid lumen when plants are transferred from light to dark, however, no experimental evidence was presented [97]. In fact, later work with thylakoid-targeted aequorin provided evidence for a dark-induced increase of [Ca^2+^] in the thylakoid lumen [24]. Two alternative chloroplast Ca^2+^ stores in the stroma have been proposed: binding to unknown protein(s) or to charges from phosphorylated proteins and phospholipids of the thylakoid membranes. This alternative scenario is unproven and unlikely, as binding to high-capacity calcium storage proteins, such as calreticulin in the ER, is accompanied by a high resting free [Ca^2+^], which conflicts with an observed resting [Ca^2+^]_str_ in the low nanomolar concentration [21]. So, the extent to which Ca^2+^ could be released from the thylakoid lumen to shape the HL-induced [Ca^2+^]_str_ dynamic remains obscure. Alternative stores such as the ER have not received much attention yet, although close contacts between ER and chloroplasts have been observed [63–65].

ER-chloroplast membrane contact sites (MCS) have long been postulated as sites of lipid exchange fueled in part by the observation of a physical association between the chloroplast and ER revealed with optical tweezers [64, 98, 99]. More recently, ER-resident proteins VESICLE ASSOCIATED PROTEIN 27-1 and −3 (VAP27-1 and −3) were postulated to be tethers of the ER-chloroplast MCS, which when mutated, led to subtle alterations in lipid composition [100]. While ER-mitochondria MCS are key players in Ca^2+^ homeostasis in mammals, involving multiple Ca^2+^ transport systems [101], a similar chloroplast-ER counterpart has yet to be identified. With that in mind, [Ca^2+^]_ER_, imaged with the ER Ca^2+^ reporter line ER-GCaMP6-210, mirrored both the fast [Ca^2+^]_str_ spike and persistent increase (Fig. 6D), which was drastically altered by CPA, a known inhibitor of type IIA Ca^2+^-ATPases (Fig. 6F).

Notwithstanding that the timescale and intensity of the HL treatment differ between our study (minutes) and that of Exposito-Rodriguez et al. (hours) [83], it is likely that the cytosolic roGFP2-Orp1 signals we detected are an effect of LOOH accumulation. In fact, regarding a potential Ca^2+^ transfer from ER to the chloroplast stroma during HL stress, the accumulation of cytosolic LOOHs is significant in connection to the recent observation that ER-mitochondrial membrane contact sites are hotspots of lipid peroxidation during ferroptosis, a form of cell death characterized by extensive lipid peroxidation [102]. Sassano et al. discovered that ferroptosis enlarges the ER-mitochondrial membrane contact site in mice fibroblasts leading to an increased ER-to-mitochondria Ca^2+^ transfer [102]. Hence, we hypothesize that ^1^O_2_-derived cytosolic LOOH modifies the ER-chloroplast contact site leading to the observed persistent [Ca^2+^]_str_ elevation, potentially through a CPA-sensitive Ca^2+^ channel or transporter (Fig. 6J). Future experiments involving roGFP2-synORP1, targeted LOOH manipulation, and mutation of Ca^2+^ transport systems will help clarify this hypothesis. At this point, we cannot exclude the accumulation of another HL-induced intermediate other than ROS that bridges photosynthesis and Ca^2+^ signaling.

## Conclusion

Inspired by the burning glass effect of water droplets on the leaf surface, we found that HL triggers a biphasic increase of [Ca^2+^]_str_ consisting of a fast Ca^2+^ peak that occurs in seemingly random chloroplasts, followed by a persistent Ca^2+^ increase in most chloroplasts. This [Ca^2+^]_str_ response is a consequence of photodamage to the electron transport chain caused by HL and the persistent increase is likely the result of ^1^O_2_ altering a Ca^2+^ transfer between the ER and the chloroplast stroma. This mechanism may help plants acclimate to HL, offering potential new strategies to enhance photosynthesis and improve crop yields. Of note, while the HL-induced persistent [Ca^2+^]_str_ increase did not resemble a heat shock-induced [Ca^2+^]_str_ response, elevated temperatures did seem to accelerate the HL response (Fig. 5). The accelerated [Ca^2+^]_str_ response could be an effect of the generally enhanced rates of biochemical reactions at higher temperatures in the range of 10°C to 40°C [78]. A combined effect of heat on the HL-induced [Ca^2+^]_str_ response could be particularly relevant in the context of future warming climates [79].

## Contributions

DK, SR, and SS designed the overall study, performed experiments and wrote the manuscript with input from SB, CS, DVD, BW, MS, MT, and AC. MS and AC provided GECIs, other genetically encoded fluorescent reporters, and expertise on sensor usage protocols. ASZ, BW, and MT cloned and produced the NTRC- and AKDE1-R-GECO1 expressing Arabidopsis lines. BMOM, SB, and AC performed the soil-grown plant confocal imaging. KZ performed the automated white light system confocal microscopy experiments, supervised by MS. EM and DVD assisted DK with temperature shift experiments. All authors gave feedback on the manuscript.

## Acknowledgements

This study was supported by the Knut and Alice Wallenberg Foundation (grant 2021.0071 to SR and SS), the European Research Council (Consolidator grant 101044878 to DK and SS), the Austrian Science Fund (Lise Meitner Program grant number M-2745-B to ASZ, and project P 37245-B to BW and MT), the EU H2020-SFS-2019-2 RIA project ADAPT (GA 2020 862-858 to BW and MT), the Ministero dell’Istruzione, dell’Università e della Ricerca Fondo per Progetti di Ricerca di Rilevante Interesse Nazionale 2022 (PRIN 2022NMSFHN) (to AC and SB) and by the Agritech National Research Center and received funding from the European Union Next-GenerationEU (PIANO NAZIO-NALE DI RIPRESA E RESILIENZA (PNRR)—MISSIONE 4 COMPONENTE 2, INVESTIMENTO 1.4–D.D. 1032 17/06/2022, CN00000022 to BMOM). Part of the work was carried out with the support of the NOLIMITS Center of Excellence for Plant Biology and Other Life Sciences established by the University of Milan. MS thanks the Deutsche Forschungsgemeinschaft (DFG) for funding (SCHW 1719/5-3, SCHW 1719/10-1, SCHW 1719/11-1) and for the infrastructure grant INST 211/903-1 FUGG for the Zeiss LSM 980 confocal microscope as operated by the Imaging Network of the University of Münster (RI_00497). KZ thanks the China Scholarship Council (202006910019) for a personal PhD studentship. Seeds for some of the reporter lines were kindly provided: R-GECO1 by Melanie Krebs (Heidelberg University, Germany) and GCaMP3 by Simon Gilroy (University of Wisconsin-Madison, USA) and Masatsugu Toyota (Saitama University, Japan).

---

**Fig. Suppl. 1.**
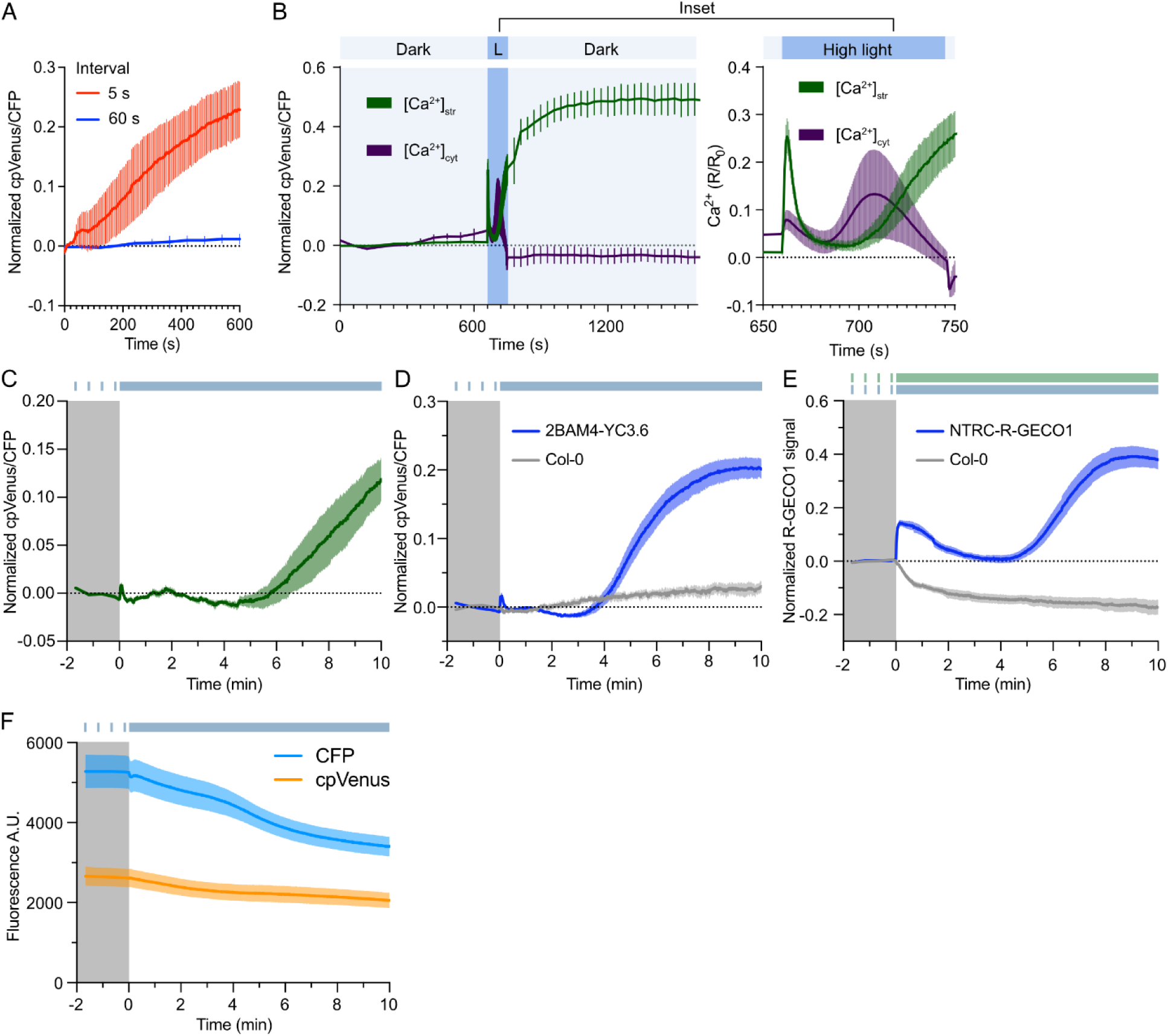
[Ca^2+^]_str_ dynamics in different light regimes and tissues; Col-0 controls for 2BAM4-YC3.6 and NTRC-R-GECO1 lines. **(A)** [Ca^2+^]_str_ dynamics induced by blue light (436 +/− 20 nm) with different intervals between blue light stimuli (5 s or 60 s) in cotyledons expressing 2BAM4-YC3.6. The blue light stimulus was present during the entire exposure time of an image (300 ms) taken on a widefield microscope. n = 3. (**B**) [Ca^2+^]_str_ and [Ca^2+^]_cyt_ dynamics induced by 90 seconds of blue light (436 +/− 20 nm) in the cotyledons of the 2BAM4-YC3.6 and NES-YC3.6 expressing lines, respectively. L (dark blue bar) is the continuous succession of 300 ms images on a widefield microscope and is enlarged in the inset. n = 3. (**C**) [Ca^2+^]_str_ dynamics induced by 10% 405 nm laser (blue bar) in the leaves of the 2BAM4-YC3.6 line. n = 5. (**D**) Autofluorescence of Col-0 cotyledons (n = 5) and signals of 2BAM4-YC3.6 cotyledons (n = 6) induced by 10% 405 nm laser (blue bar) using the same imaging settings. (**E**) Autofluorescence of Col-0 cotyledons (n = 5) and signals of NTRC-R-GECO1 expressing cotyledons (n = 5) induced by 10% 405 nm laser (blue bar) and 1% 561 nm laser (green bar) using the same imaging settings. (**F**) cpVenus and CFP fluorescence of 2BAM4-YC3.6 cotyledons (n = 6) induced by 10% 405 nm laser (blue bar).

**Fig. Suppl. 2.**
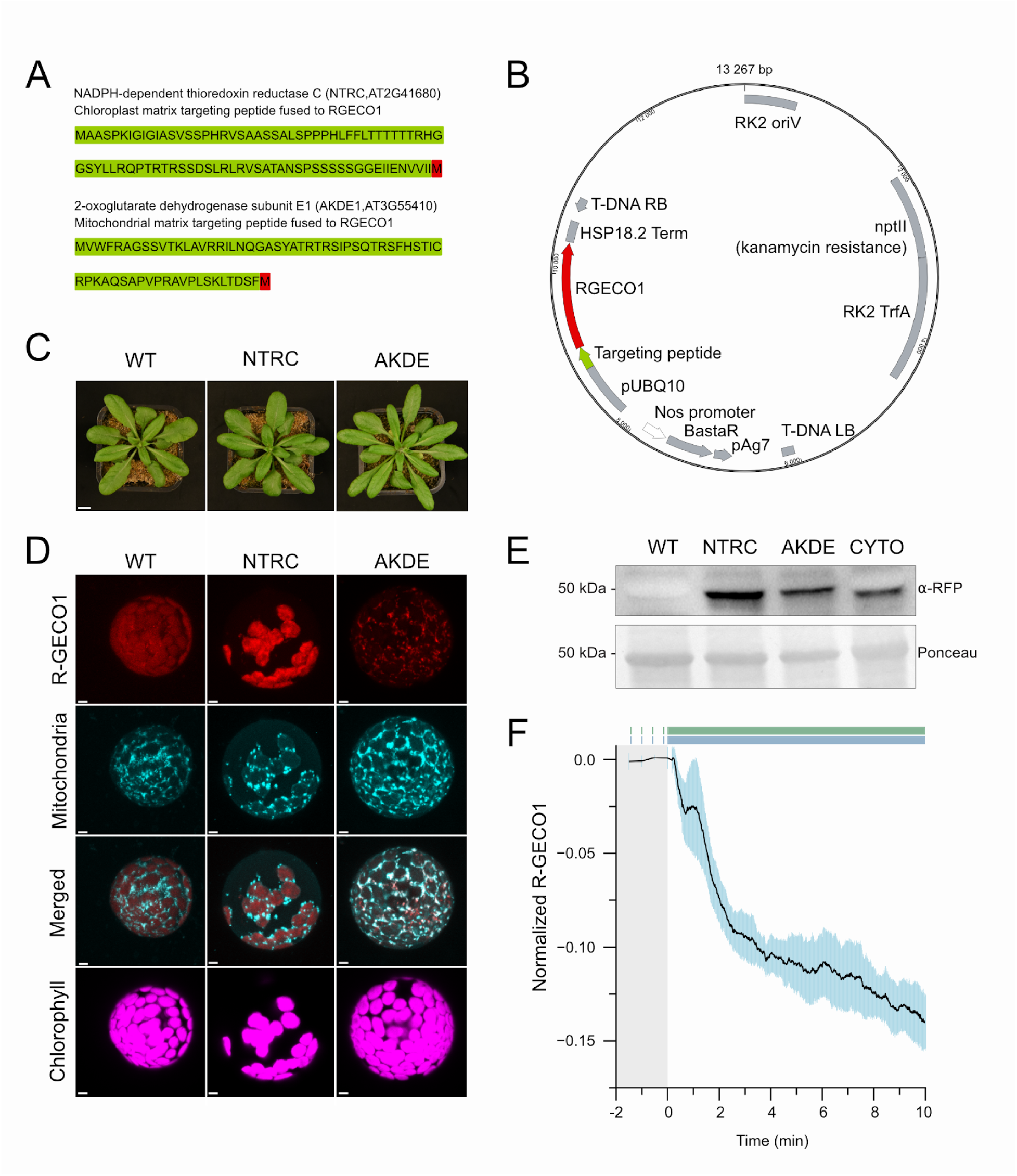
Characterisation of the newly established NTRC- and AKDE1-R-GECO1 lines. **(A)** Amino acid sequences of the NTRC (chloroplast) and AKDE1 (mitochondria) targeting peptides that were used to target the sensors. **(B)** Plasmid map of the vector used to transform *Arabidopsis thaliana* (ecotype Col-0). Expression of the GECIs is driven by an Arabidopsis UBIQUITIN10 (UBQ10) promotor. The structure of both plasmids is the same, except for the sequences of the targeting peptides (NTRC or AKDE). **(C)** Representative images of mature NTRC- and AKDE1-R-GECO1 Arabidopsis rosettes compared to wild type (WT). Scale bar, 1 cm. **(D)** Localization of NTRC- and AKDE1-R-GECO1 in leaf mesophyll protoplasts derived from mature leaves. Mitochondria were stained with a MitoView™ 405 dye. Scale bar, 5 μm. **(E)** Western blot on protein extracts from wild type (WT), NTRC- and AKDE1-R-GECO1, and a previously established cytosolic R-GECO1 (CYTO) line with anti-RFP. Ponceau S stain of the Rubisco large subunit as the protein serves as loading control. **(F)** [Ca^2+^]_str_ dynamics induced by 10% 405 nm laser in cotyledons of AKDE-R-GECO1 expressing plants. AKDE-R-GECO1 was additionally excited by a 1% 561 nm laser (green bar). n = 3.

**Fig. Suppl. 3.**
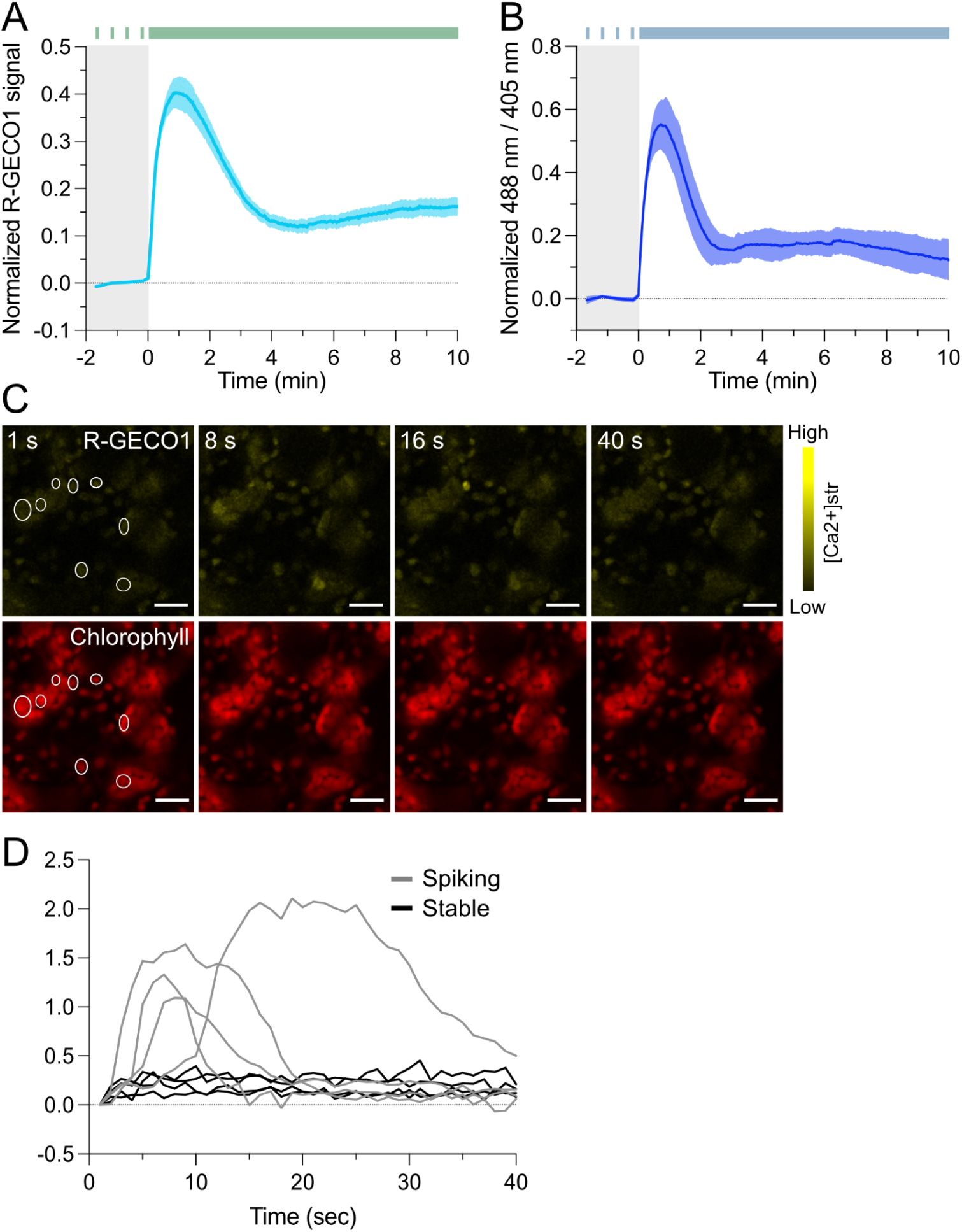
pH effects and single chloroplast [Ca^2+^]_str_ dynamics in the NTRC-R-GECO1 expressing plants. **(A)** R-GECO1 signal induced by 1% 561 nm laser (green bar) in the NTRC-R-GECO1 line. n = 4. (**B**) pH dynamics in chloroplast stroma sequentially induced by 10% 405 nm and 15% 488 nm laser (blue bar) in the cp-cpYFP line. n = 5. (**C**) R-GECO1 signal (top) and chlorophyll (bottom) at 1s, 8s, 16s and 40s onset of 10% 405 nm laser. White circles indicate single chloroplasts selected for quantification. Scale, 20 μm. (**D**) Quantified data of selected single chloroplasts, spiking ones are marked blue, while stable ones are marked red.

**Fig. Suppl. 4.**
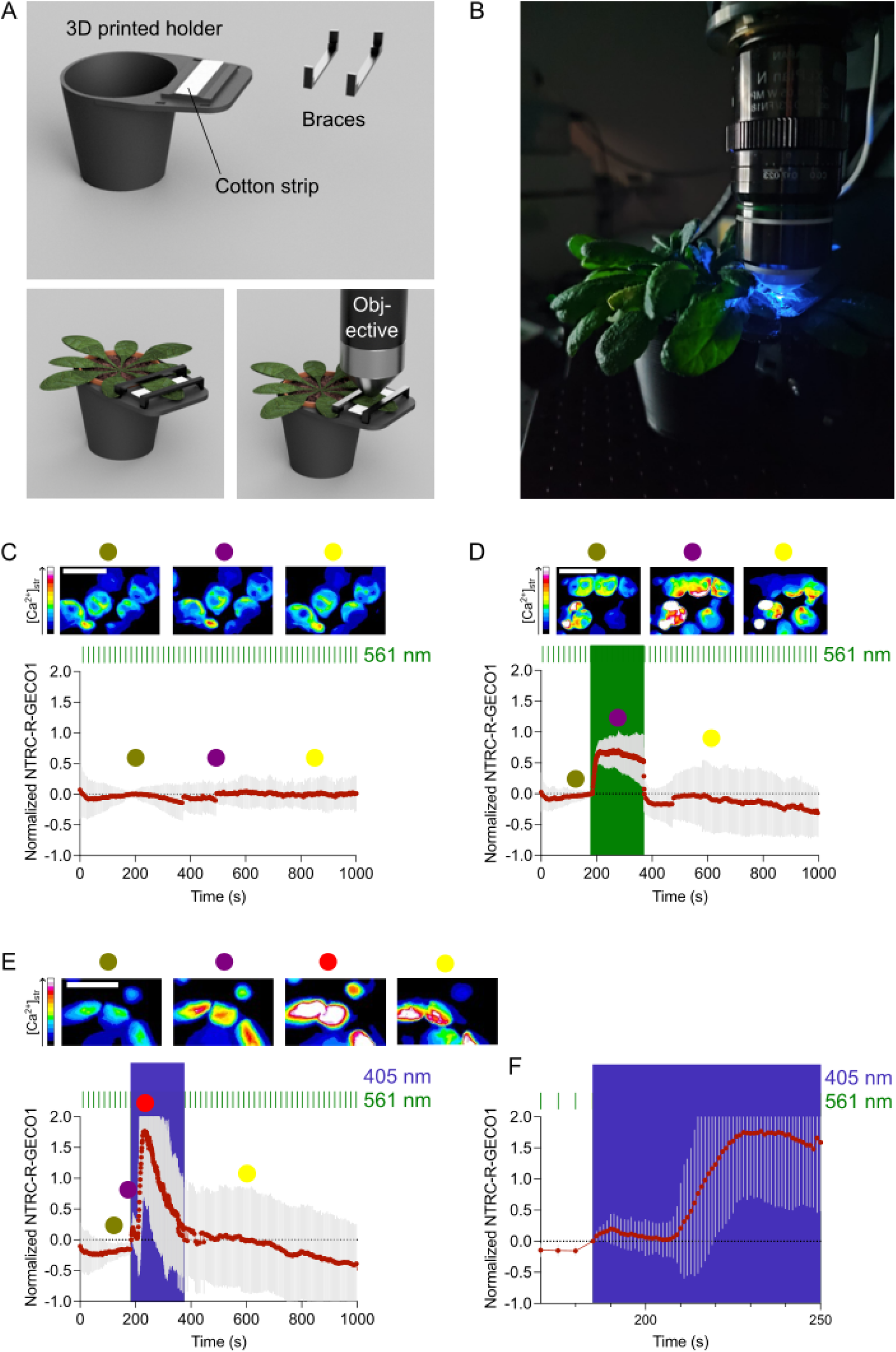
*In vivo* Confocal Laser Scanning Microscopy of chloroplasts from mature *Arabidopsis thaliana* leaves. **(A)** Model of a 3D-printed microscope holder for a soil-grown Arabidopsis plant. **(B)** Picture of the system to illustrate the setup for *in vivo* Confocal Laser Scanning Microscopy. **(C)** Control experiment for [Ca^2+^]_str_ dynamics. NTRC-R-GECO1 expressing plant was illuminated with 1% 561 nm laser every 5 seconds for 16 minutes. Average of 40 individual chloroplasts from 5 experiments (n). **(D)** NTRC-R-GECO1 expressing plant was illuminated with 1% 561 nm laser every 5 s for 3 min, followed by 3 min of continuous illumination, and then 10 min of illumination and acquisition every 5 s. Average of 40 individual chloroplasts from 4 experiments (n). **(E)** NTRC-R-GECO1 expressing plant was illuminated with 1% 561 nm laser every 5 s for 3 min, followed by 3 min of continuous illumination with 1% 561 laser and 20% 405 nm illumination, and then 10 min of illumination with 1% 561 nm laser and acquisition every 5 s. Inset images of single chloroplasts. Average of 50 individual chloroplasts from 5 experiments (n). **(F)** Inset of the blue light induced [Ca^2+^]_str_ dynamic in (E).

**Table Suppl. 1:** Overview of laser intensities. Power at the focal point was measured on the different microscopes with a Microscope Slide Power Meter (Thorlabs; in µWatt, bold text) and converted to PAR (μmol photons m^−2^ s^−1^).

| $\mu\text{Watt} / \mu\text{mol m}^{-2} \text{s}^{-1}$ | | Laser strength | | | | | | | | |
| --- | --- | --- | --- | --- | --- | --- | --- | --- | --- | --- |
|  |  | 1% | 2% | 5% | 7.5% | 10% | 15% | 20% | 30% | 50% |
| Zeiss LSM 780 | 405 nm | **10,8** / 202,4 | **22** / 412,3 | **56** / 1049,5 | **86** / 1611,7 | **113** / 2117,7 | **175** / 3279,7 | **235** / 4404,1 | - | **580** / 10869,8 |
|  | 488 nm | - | - | - | - | - | - | **110** / 2484,0 | - | - |
|  | 561 nm | **24** / 623 | **47** / 1220,1 | **120** / 3115,2 | - | **240** / 6230,3 | **355** / 9215,7 | **457** / 11863,6 | - | **1200** / 31151,7 |
|  | 633 nm | **5,8** / 169,9 | **11,8** / 345,6 | **29,5** / 864,1 | - | **59** / 1728,2 | **92,5** / 2709,5 | **124** / 3632,1 | **180** / 5280,8 | **310** / 9080,4 |
| PerkinElmer (spinning disk) | 405 nm | - | - | **78,07** / 2191,5 | - | **149** / 4182,5 | **220** / 6175,6 | **297** / 8337 | - | - |
| Nikon Eclipse Fn (Nikon A1R confocal) | 405nm | - | - | - | - | - | - | **120** | - | - |
|  | 561 nm | **26** | - | - | - | - | - | - | - | - |

**Fig. Suppl. 5.**
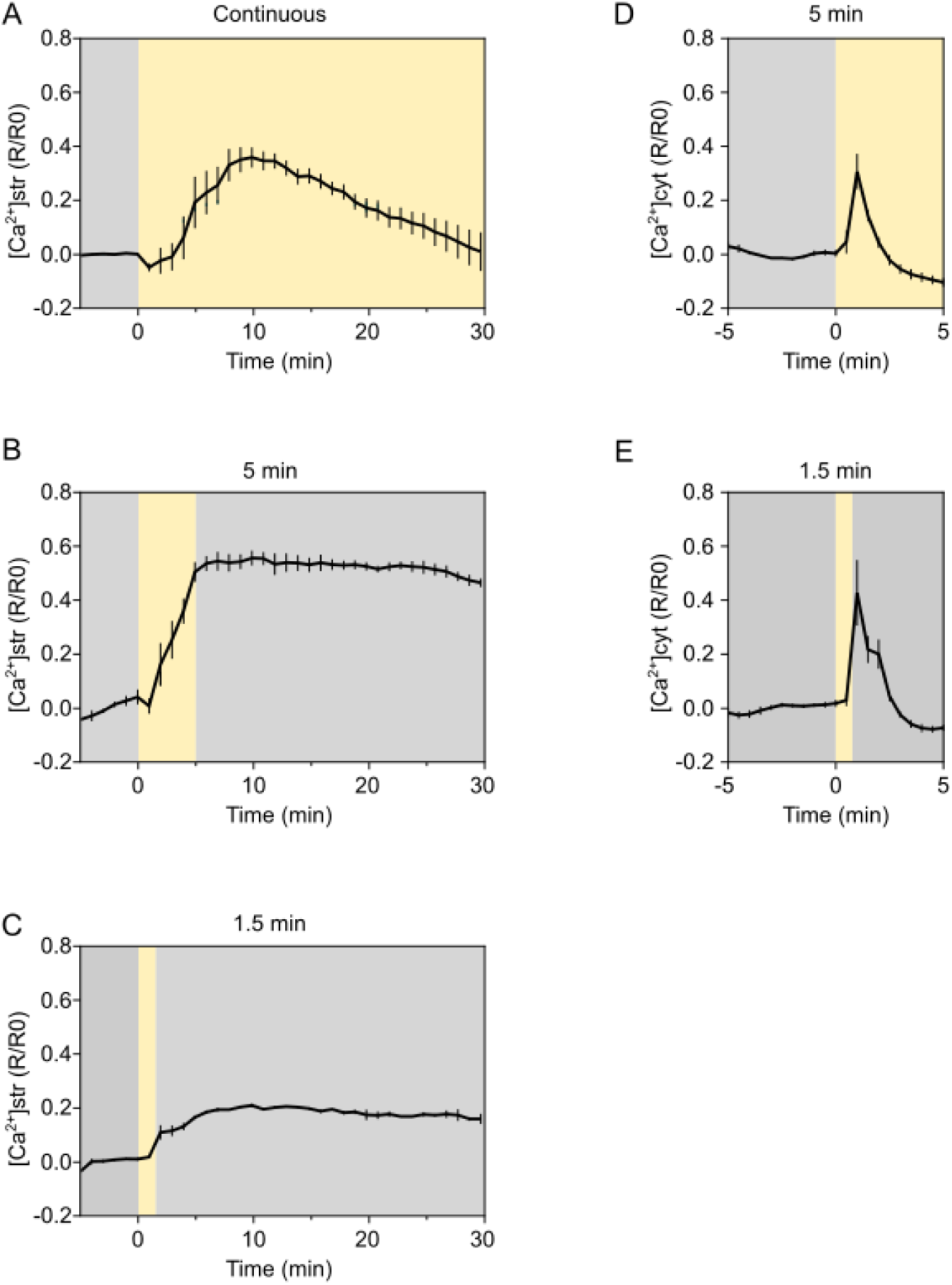
White light-induced [Ca^2+^]_str_ and [Ca^2+^]_cyt_ responses. **(A-C)** [Ca^2+^]_str_ dynamics induced by a white halogen light source in cotyledons of plants expressing 2BAM4-YC3.6. Light treatments (continuous, 5, or 1.5 min) are indicated with a yellow bar. Cotyledons were intermittently imaged every 1 min, indicated by the error bars (SEM), between the white light stimuli. **(A)** Continuous (n = 7), **(B)** 5 min (n = 3), or **(C)** 1.5 min (n = 2). **(D-E)** [Ca^2+^]_cyt_ dynamics induced by a white halogen light source in cotyledons of the NES-YC3.6 expressing plants. Cotyledons were intermittently imaged every 30 s, indicated by the error bars (± SEM), between the white light stimuli. **(D)** 5 min (n = 4), or **(E)** 1.5 min (n = 6). YC3.6 ratio divided by the average of the ratios before stimulus (R/R0).

**Fig. Suppl. 6.**
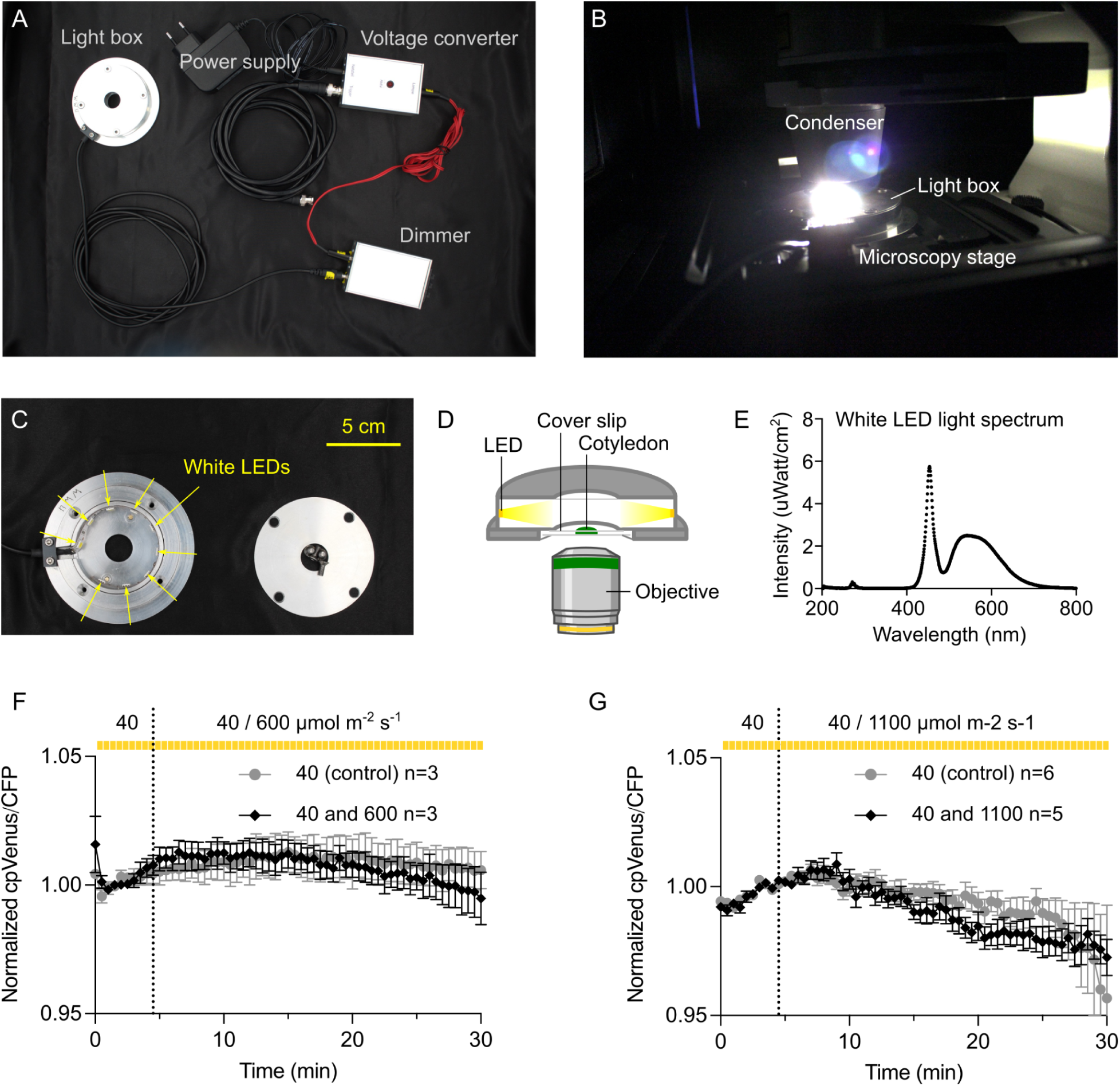
An automated illumination system with white light does not trigger [Ca^2+^]_str_ dynamics. **(A)** Overview of equipment for the automated illumination system. **(B)** Position of the light box on the microscope. **(C)** Detail of the inside of the light box with white light LED strip. **(D)** Diagram of the light box setup and position of the sample. **(E)** Recorded light spectrum of the white LEDs. **(F-G)** [Ca^2+^]_str_ dynamics in presence of white LED light in cotyledons of 2BAM4-YC3.6 expressing plants when switching from 40 to 600 μmol photons m^−2^ s^−1^ (F; n = 3), or 40 to 1100 μmol photons m^−2^ s^−1^ (G; n = 5-6) after approximately 4.5 min (time point indicated by the dotted line). Light treatment is indicated with a yellow bar. Cotyledons were intermittently imaged every 30 s between the white light stimuli.

**Fig. Suppl. 7.**
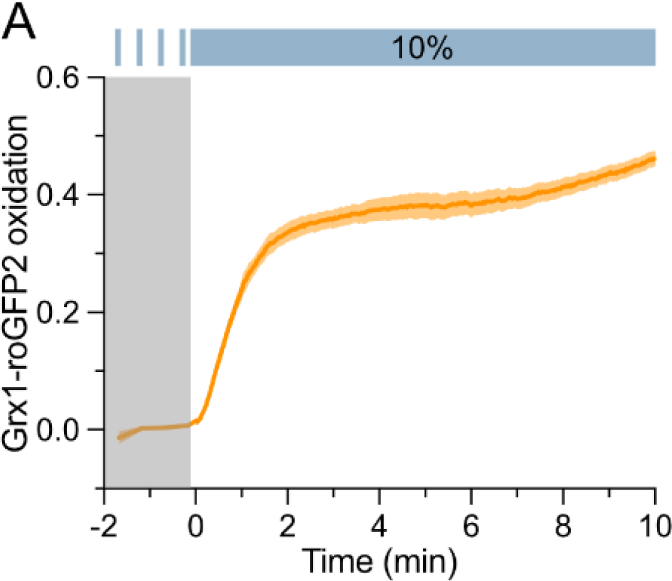
Stromal glutathione redox potential upon blue light stimulus. **(A)** Stromal ROS induced by 10% 405 nm laser (blue bar) in cotyledons of TKTP-Grx1-roGFP2 expressing plants. n = 5.

**Fig. Suppl. 8.**
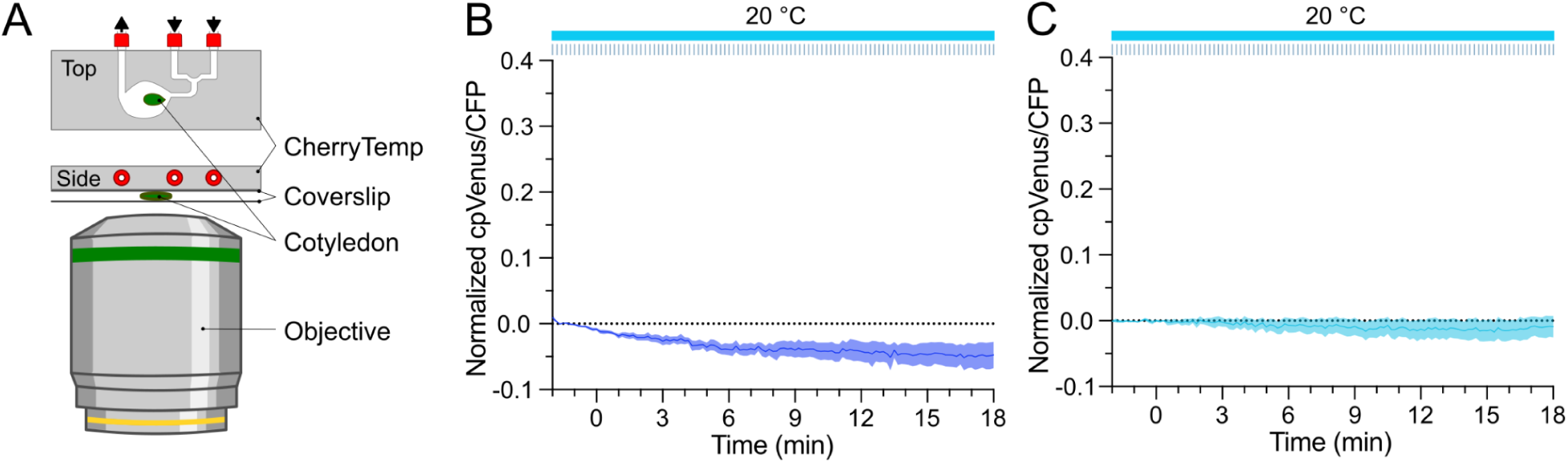
Imaging setup and controls for temperature shift. **(A)** A diagram of imaging setup with temperature control based on microfluidics heating and cooling of a chip (CherryTemp heating/cooling system top and side view). (**B**) [Ca^2+^]_str_ dynamics of cotyledons expressing 2BAM4-YC3.6 under the same light regime of Fig. 4B, at a constant temperature of 20℃, n = 5. (**C**) [Ca^2+^]_str_ dynamics of roots of 2BAM4-YC3.6 under the same light regime of Fig. 4B, at a constant temperature of 20℃, n = 5.

**Fig. Suppl. 9.**
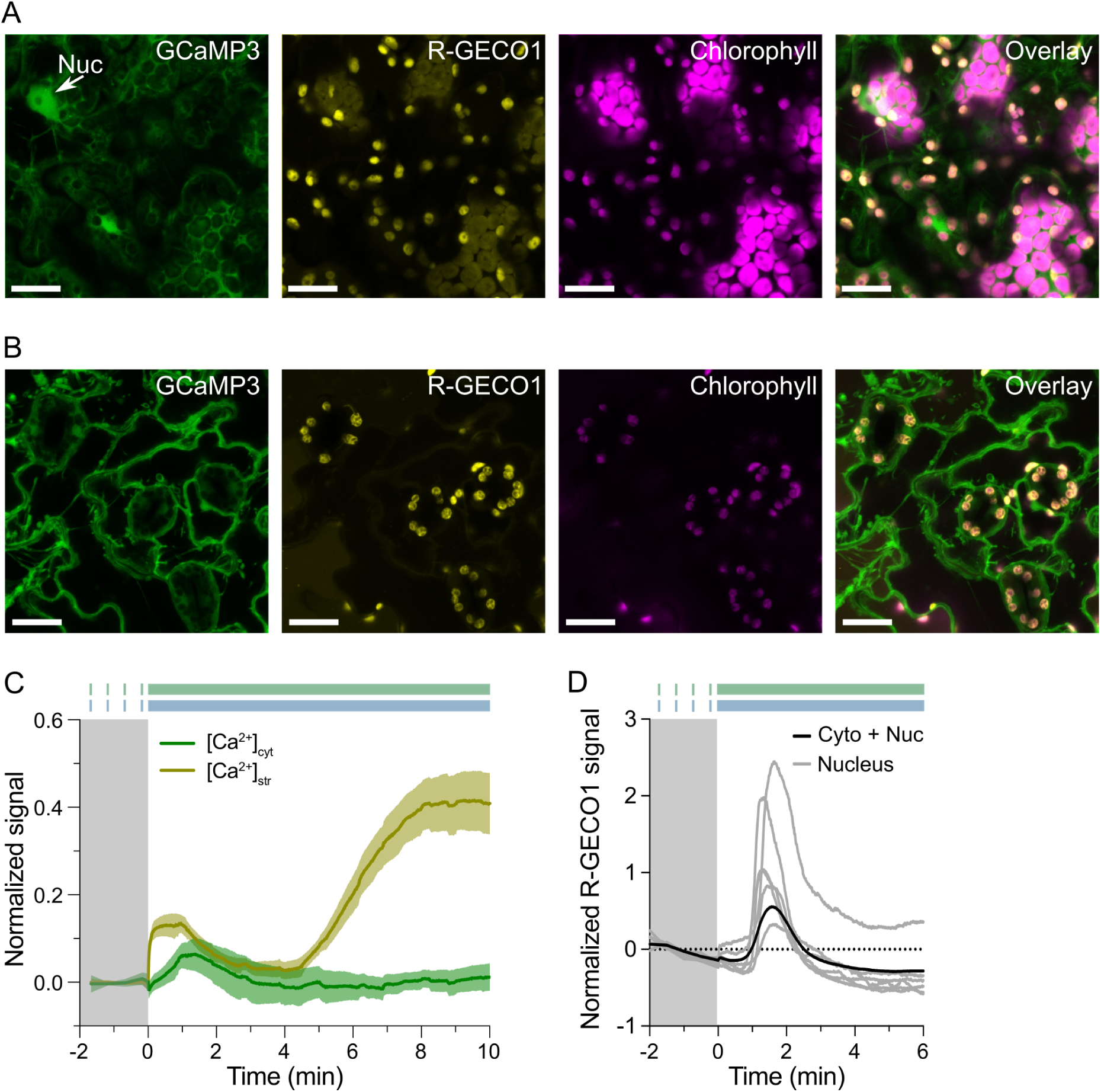
[Ca^2+^] dynamics in cytosol and stroma in NTRC-R-GECO1 × cyto-GCaMP3. **(A)** GCaMP3, R-GECO1 and chlorophyll signal in epidermal/mesophyll cells of cotyledons of NTRC-R-GECO1 × cyto-GCaMP3 dual marker lines. GCaMP3 localizes to the cytosol and to the nucleus (indicated with an arrow; Nuc). (**B**) GCaMP3, R-GECO1 and chlorophyll signal in guard cells of cotyledons of NTRC-R-GECO1 × cyto-GCaMP3 expressing plants. Scale bar, 20 μm. (**C**) [Ca^2+^]_cyt_ and [Ca^2+^]_str_ dynamics induced by 10% 405 nm laser light (blue bar) in cotyledons of the NTRC-R-GECO1 × cyto-GCaMP3 dual marker line, NTRC-R-GECO1 was additionally excited by 1% 561 nm laser (green bar). n = 5. **(D)** [Ca^2+^] dynamics induced by 20% 405 nm laser in selected ROIs of nuclei compared to the combination of cytosol and nucleus (quantification of full field of view of the image) in a cyto-nuclear localized R-GECO1-expressing line. Cyt-R-GECO1 was additionally excited by 5% 561 nm laser (green bar). Quantification of one representative experiment shown in Suppl. Movie. 4.

**Fig. Suppl. 10.**
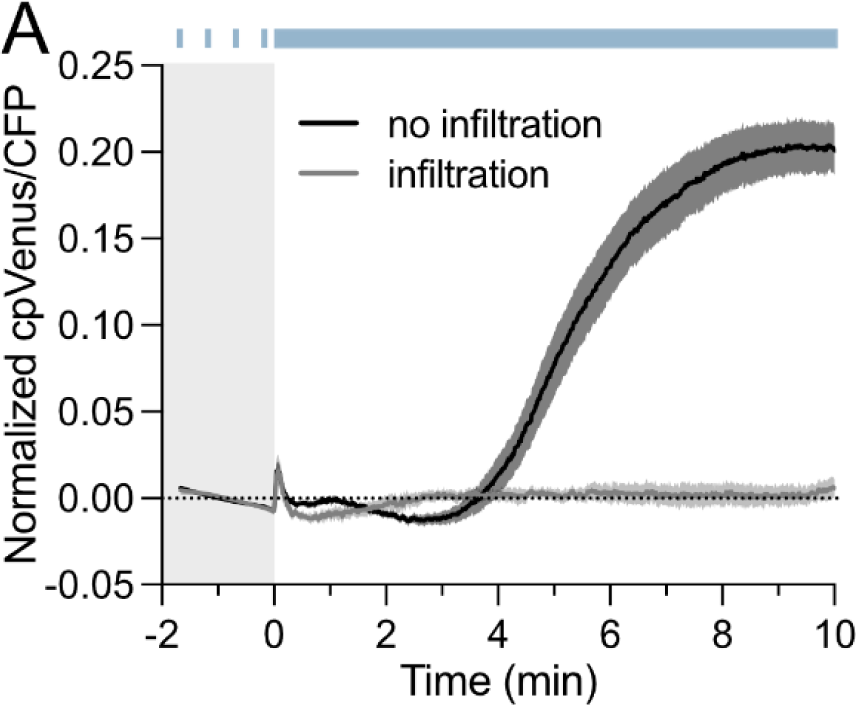
HL-induced [Ca^2+^]_str_ with or without water infiltration. **(A)** [Ca^2+^]_str_ dynamics induced by 10% 405 nm laser (blue bar) in cotyledons of 2BAM4-YC3.6 expressing plants, with water infiltration (n = 5) or without water infiltration (n = 6).

**Fig. Suppl. 11.**
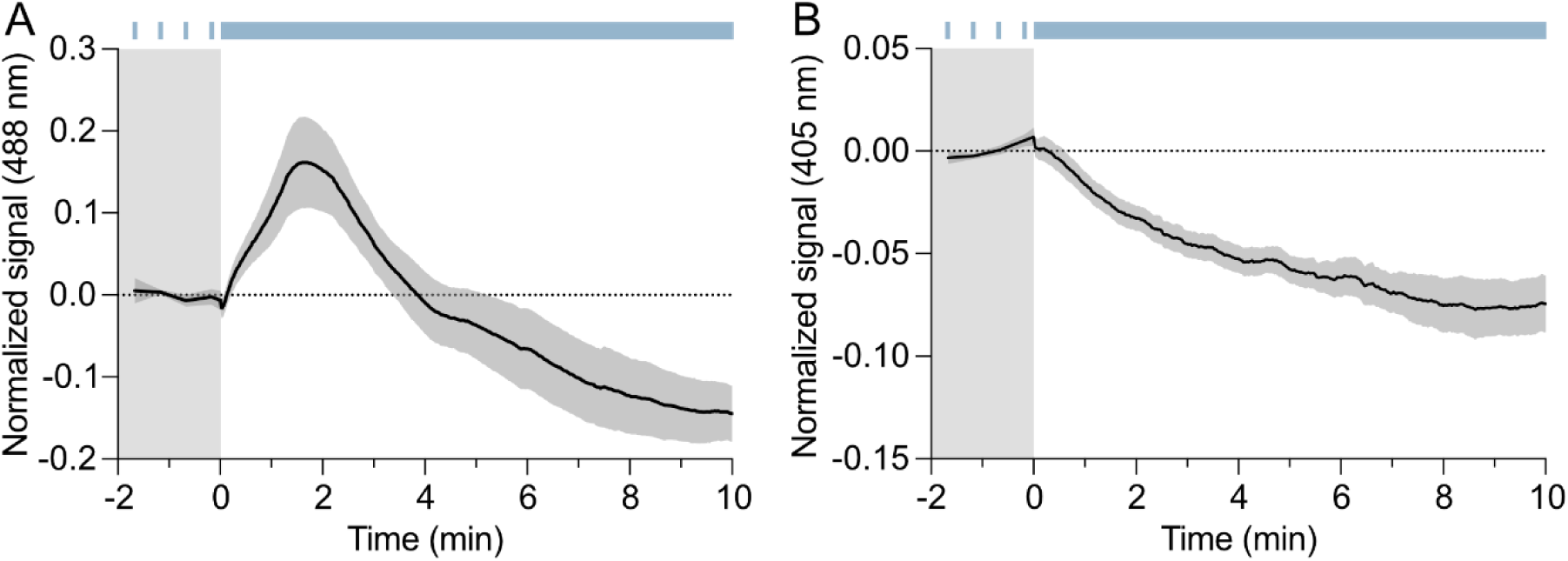
Normalized signal of ER-GCaMP6-210 excited by 488 nm or 405 nm. **(A)** Normalized signal excited by 488 nm laser upon a HL trigger of sequential 10% 405 nm / 20% 488 nm laser (blue bar) in cotyledons of ER-GCaMP6-210 expressing plants (n = 6). (**B**) Normalized signal excited by 405 nm laser upon a HL trigger of sequential 10% 405 nm / 20% 488 nm laser (blue bar) in cotyledons of ER-GCaMP6-210 expressing plants (n = 6).

